# Estimates for the astrocyte endfoot sheath permeability of the extra-cellular pathway

**DOI:** 10.1101/2022.11.16.516727

**Authors:** Timo Koch, Vegard Vinje, Kent-André Mardal

## Abstract

**Background:** Astrocyte endfoot processes are believed to cover all micro-vessels in the brain cortex and may play a significant role in fluid and substance transport into and out of the brain parenchyma. Detailed fluid mechanical models of diffusive and advective transport in the brain are promising tools to investigate theories of transport.

**Methods:** We derive theoretical estimates of astrocyte endfoot sheath permeability for advective and diffusive transport and its variation in microvascular networks from mouse brain cortex. The networks are based on recently published experimental data and generated endfoot patterns are based on Voronoi tessellations of the perivascular surface. We estimate corrections for projection errors in previously published data.

**Results:** We provide structural-functional relationships between vessel radius and resistance that can be directly used in flow and transport simulations. We estimate endfoot sheath filtration coefficients in the range *L*_*p*_ = 0.2 × 10^−10^ m Pa^−1^ s^−1^ to 2.7 × 10^−10^ m Pa^−1^ s^−1^, diffusion membrane coefficients in the range *C*_*M*_ = 0.5 × 10^3^ m^−1^ to 6 × 10^3^ m^−1^, and gap area fractions in the range 0.2 % to 0.6 %.

**Conclusions:** The astrocyte endfoot sheath surrounding microvessels forms a secondary barrier to extra-cellular transport, separating the extra-cellular space of the parenchyma and the perivascular space outside the endothelial layer. The filtration and membrane diffusion coefficients of the endfoot sheath are estimated to be an order of magnitude lower than the extra-cellular matrix while being two orders of magnitude higher than the vessel wall.

## Background

Astrocyte endfoot processes have been reported to cover virtually all of the microvasculature in brain gray matter [1, 2, 3, 4, 5, 6]. The endfoot processes overlap [3] and form a sheath that constitutes the outer boundary of the perivascular space (PVS). Exchange of fluid across the endfoot sheath is vital to maintain homeostasis of the central nervous system [7], and a key component of the proposed glymphatic theory [8]. The extra-cellular transport pathway, between individual endfoot processes, and the associated permeabilities of the endfoot sheath are relevant for the interpretation of transport phenomena observed for cerebrospinal fluid (CSF), and passively transported substances that are believed to not enter astrocytes in large quantities (such as many MRI contrast agents) and are used in the analysis of flow and transport processes into, out of and within the brain parenchyma. We omit here the discussion of intra-cellular pathways (see [8, 9, 10, 11] for proposed roles and scientific debate) and perivascular pathways (see e.g. [6]).

The astrocyte endfoot sheath enclosing the microvessels in brain tissue can be viewed as the surface of a tube tiled by individual endfoot processes, cf. [6, Fig.2]. Voronoi tessellations have been successfully used to describe the geometric configuration of cell populations and cell dynamics for decades [12]. Voronoi tessellations appear if cells are grown radially from a center point at constant speed until collision with a neighbor cell growing at the same speed, a simulation process used by [13] to construct virtual astrocyte endfoot processes. However, Voronoi tessellations of a point set can also be more directly constructed as the dual graph of a Delaunay triangulation of the point set. Motivated by the recent work of Wang et al. [6], in which the authors visualized endfoot process gaps in mouse brain resembling Voronoi tessellations, we herein propose their use to generate artificial endfoot patterns. An exemplary realization of such a pattern is shown in Fig. 1 (cf. [6, Fig.2]). To estimate the permeability of the generated cell patterns, the parameterized surface model representation has to be combined with a corresponding cross-sectional model representation. A schematic cross-sectional cut through a capillary in Fig. 2 introduces the considered perivascular structures and parameterization of the interendfoot gaps.

**Figure 1.**
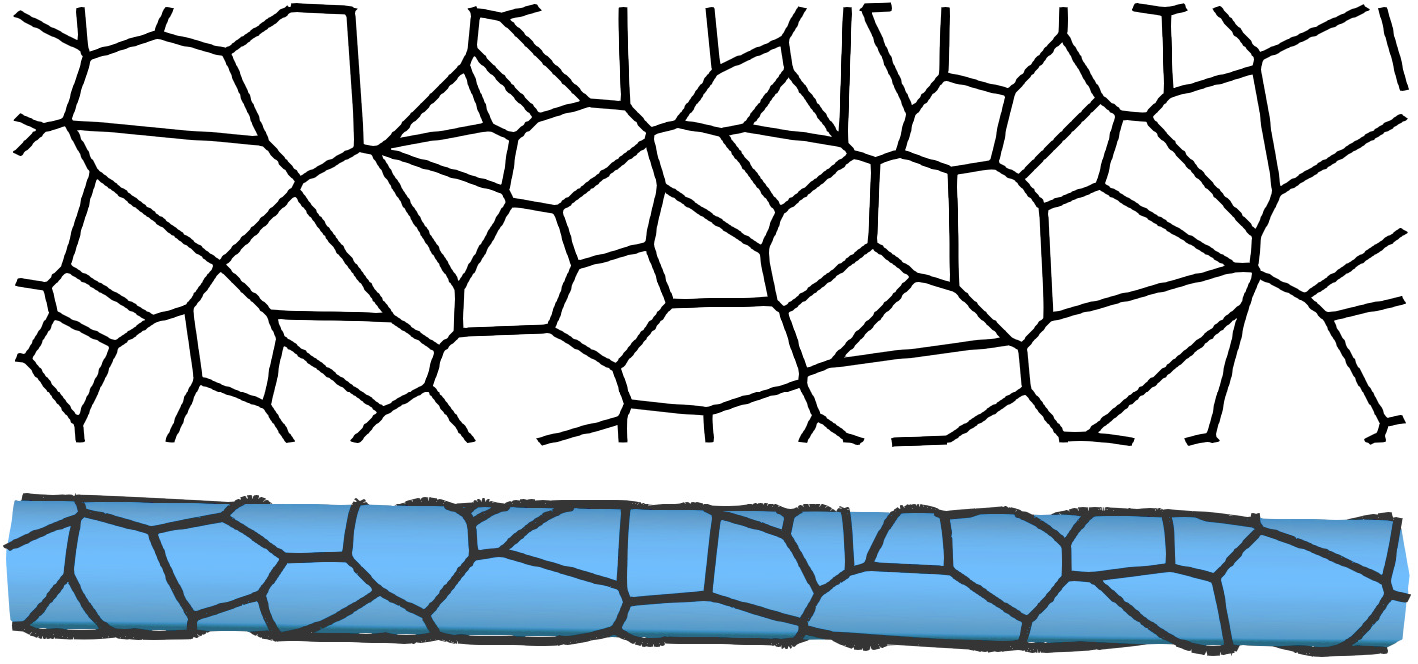
Voronoi tessellation model of endfoot sheath. Top: a sample Voronoi tessellation with a mean endfoot area of *A* = 25 µm^2^ and a vessel radius of *r*_o_ = 10 µm^2^ (including endfoot sheath). Cell center positions are generated from a uniform random distribution. The tessellation is periodic which is important to ensure that wrapping around a vessel will provide a consistent tiling of the vessel surface. Bottom: same tessellation as above, but mapped onto a cylinder surface (vessel surface).

**Figure 2.**
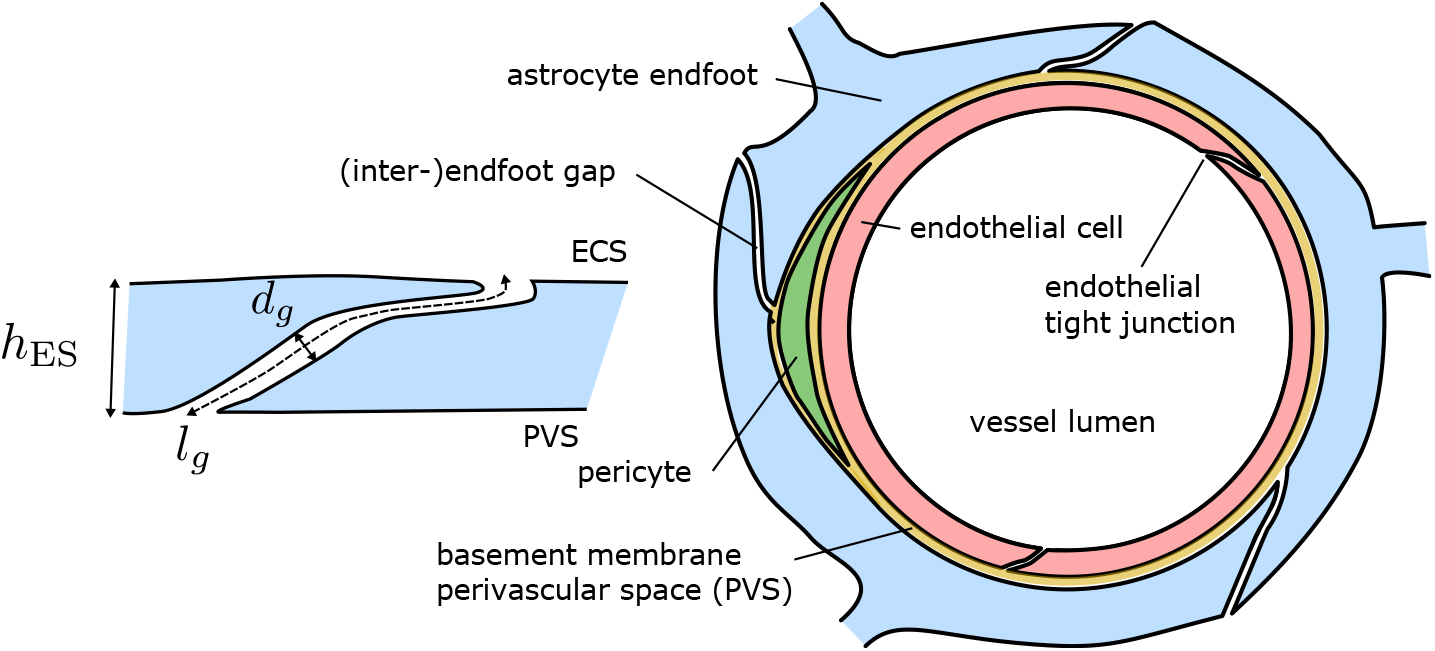
Schematic cross-section through a cortical capillary. Elements are not drawn to scale. Left: schematic of an inter-endfoot gap showing overlap between neighboring cells and identifies symbols for some geometrical measures used in this work. Right: schematic cross-sectional cut through a capillary. The ‘gaps/ring’ count for the shown cross-section is 3. The gap area fraction is measured on the outer endfoot surface (side of the extra-cellular space, ECS). For larger parenchymal microvessels (arterioles, venules), the perivascular space contains other pericytes such as smooth muscle cells and is thicker. This work is concerned with estimating the permeability of the blue astrocytic endfoot process layer relevant for fluid and substance exchange between PVS and ECS.

Few quantitive experimental studies have been published on the geometrical configuration of astrocyte endfeet. Mathiisen et al. [3] conducted an ultra-structural analysis of astrocyte endfeet in capillary vessels in rats, reporting on gap sizes, thickness, and coverage. Individual endfoot processes are separated from neighboring endfeet by gaps of 20 nm on average [3]. Moreover, neighboring endfeet are overlapping [3] and are regularly connected at gap junctions as narrow as 5 nm as described in an early ultrastructural analysis by Brightman and Reese [14]. McCaslin and coworkers [5] report average endfoot densities and average endfoot sheath thickness for capillaries and larger arterial and venous vessels using two-photon microscopy acquired in-vivo in mice. Recently, Wang et al. [6] analyzed variations in astrocytic endfoot sizes along the vascular tree in mouse brain cortex and hippocampus exvivo using confocal microscopy and their data demonstrates significant differences in endfoot sizes between venous and arterial vessels.

Permeability estimates of the endfoot sheath and its variance in microvascular networks are crucial parameters for computational models of diffusive and advective transport in brain tissue [15, 16]. Previous estimates of the permeability of perivascular and interstitial compartments have been obtained in a number of works [9, 17, 16, 18]. Asgari et al. [9] estimated the resistance of astrocyte inter-endfoot gaps based on an idealized geometrical configuration. In [18], this estimate was extended to obtain a brain-wide resistance between the periarterial/perivenous compartments and the extra-cellular space.

There are two shortcomings of the previous analyses. Firstly, the available data has only been partially combined into permeability estimates. As permeability is a crucial material property, we here aim to provide a derivation and resulting estimates. Secondly, the variation of permeability values within cortical microvascular networks has not been estimated. For example, Mathiisen et al. [3] estimated the (inter-endfoot) gap area fraction based on cross-sectional data. Wang et al. [6] reported variations of the average endfoot vessel coverage area with vessel type, and estimated resulting water flux variations, but no resulting gap area fractions. That means the data cannot be directly compared and mapping variation onto a microvascular network requires additional data or model assumptions.

In this work, we will focus on the estimation of the extra-cellular endfoot sheath permeability and its variability incorporating data on endfoot sheath ultra-structure, endfoot surface area, and variations with vessel diameters. To this end, we estimate parameter distribution in microvascular networks extracted from mouse brain [19]. Based on and parameterized by values from published experimental data [3, 6], we propose a theoretical model based on random tessellations of the endfoot sheath. The model provides estimates for the permeability of the astrocyte endfoot sheath around microvessels to transmembrane transport of fluids and transported substances. By use of this model, we can connect and compare the data obtained by Mathiisen et al. [3] (gap area fraction) and Wang et al. [6] (area variation) and discuss both permeability variations within a microvascular network and network-averaged quantities.

## Methods

### Theoretical model of endfoot sheath cell areas and gaps

To generate artificial endfoot sheath coverage patterns, we drew uniformly distributed random points on the endfoot sheath surface^[1]^. Next, we computed a Voronoi tessellation of the point set^[2]^. The tessellations consist of polygonal faces. Each polygon represents the visible surface^[3]^ covered by an endfoot process and the polygon edges (also called bisector edges) mark the location of endfoot-endfoot gaps. The surface is represented by a rectangle of width 2*πr*_o_ (where *r*_o_ is the end-foot sheath radius) and height *L* such that the total area divided by the number of polygons equals the desired mean endfoot area, *A*^[4]^. Based on the reported image data by Wang et al. [6], we assumed that the reported vessel diameters include the endfoot sheath.

Assuming a constant gap width *d*_g_, we computed *ϕ*_g_, the area fraction of the surface occupied by inter-cellular gaps (i.e. the surface available for transmembrane exchange via the extra-cellular pathway). For this, we multiplied the total edge length *l*_Σ_ in the Voronoi tessellation with the gap width and divide by the total surface area, *ϕ*_g_ := *dl*_Σ_*/*(*L*2*πr*_o_). Since the estimated gap area fractions are below 1 %, we neglect the influence of considering finite-sized gaps on the endfoot area.

For comparison with previously published data, we also computed the average number of gaps counted in cross-sectional cuts through the vessel as ‘gaps/ring’:=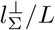, where 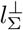 is the total length of the edges after projecting each edge in axial vessel direction. A related number is 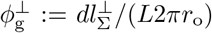 corresponding to the (reduced) gap area fraction obtained when using ‘gaps/ring’ as the basis for its estimation (as for example used in [18]).

### Combination with experimental data

Wang et al. [6] describe how the area covered by single endfoot process varies along the vascular tree for vessels of different diameters. They report endfoot areas for all analyzed vessels [6, Fig.2] and endfoot areas separate for arterial vessel and venous vessel for vessels with radius *r*_o_ > 7.5 µm [6, Fig.4]. To extract the data shown by Wang et al. [6], we used the open-source image analysis tool WebPlotDigitizer [21]. We extracted the linear regression curve (vessel average) of endfoot area as a function of vessel diameter, and all individual data points and regression trends of the data were classified into arteries and veins. To obtain an estimate over the whole range of vessels, separated into arterial and venous vessels, we constructed functions to fit well the entire range of diameters reported by Wang et al. [6]. The linear regression trends of [6] and our approximation overlaid are shown in Fig. 3.

**Figure 3.**
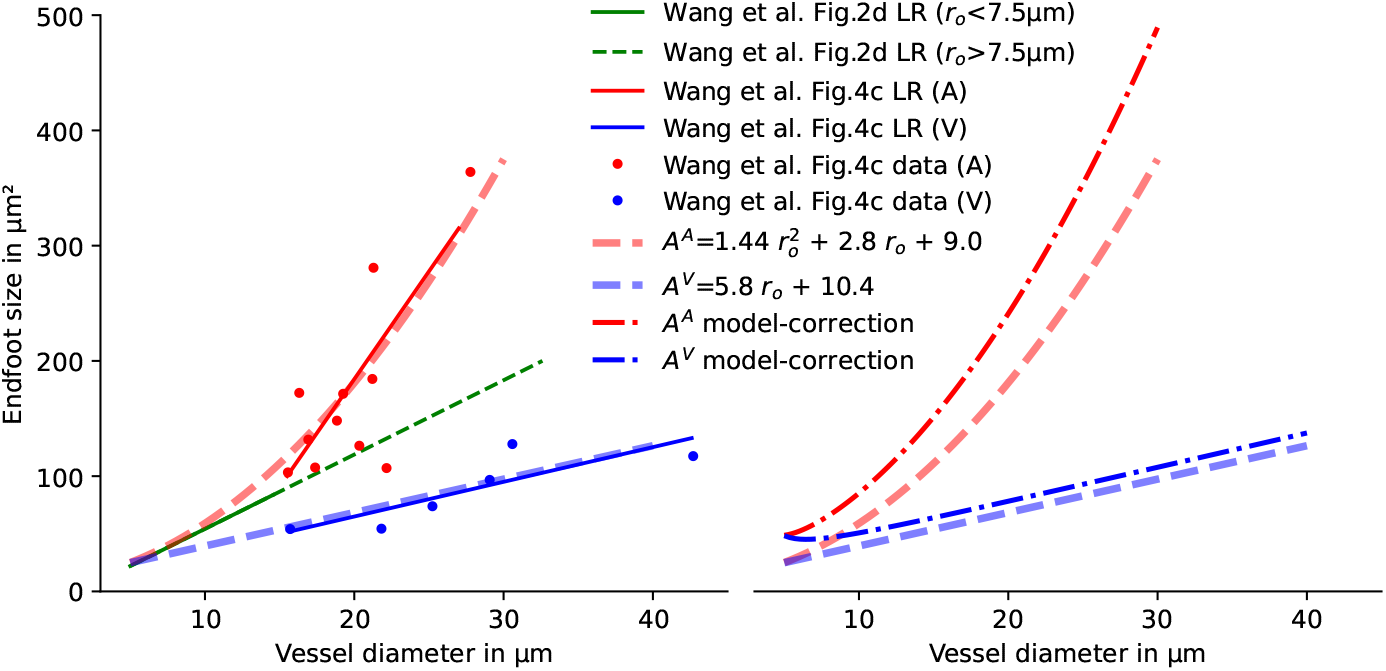
Functional relation between vessel diameter and endfoot area. Left: the proposed functional relations are based on quadratic (arteries, A) and linear (veins, V) functions such that the linear regression (LR) results from [6] are well-matched over the whole range of vessel diameters differentiating between arterial and venous vessels. The data from [6, Fig.2d, green solid line] corresponds to the data from 15-month-old mice; the data from [6, Fig.4c] (red and blue) to data from 12-month-old mice. No significant age-dependence of the endfoot area is observed by Wang et al. [6]. Dots show vessel-averaged data reported by Wang et al. [6, Fig.4c]. Average endfoot sizes, *A*^*A*^ and *A*^*V*^, for the endfoot sheath around arterial and venous vessels, respectively, are given in µm^2^ for vessel radius (including endfoot sheath) *r*_o_ in µm. Right: average endfoot sizes, *A*^*A*^ and *A*^*V*^ estimated from the data [6] and corrected relation (computed with the presented theoretical model) accounting for the error inherent to the 2D image analysis.

The areas measured by [6] correspond to plane projections of the endfoot area resulting from the analysis of 2D images rather than 3D reconstructions of the endfoot sheath, cf. [6, Fig.2]. The projection into the image plane underestimates the actual endfoot area by introducing two sources of error: (1) orthogonal projection distorts the vessel surface, and (2) half of the vessel surface is not visible in the projection. Both effects are stronger for smaller vessels where endfeet typically wrap around the vessel. We first quantified these errors based on the generated Voronoi tessellations and virtual projection as described in more detail in Appendix A. We then found a unique mapping between measured and corrected areas, which allows us to correct the projection error, see Appendix A. The second graph in Fig. 3 shows the diameter-area relationship after correction predicted by the model. This corrected diameter-area relationship is used as the basis for all parameter estimates in this work.

### Permeability for diffusive transport of passive tracers

As proposed previously, e.g. [9, 18], we conceptually model the endfoot sheath as a porous medium. Since we here only consider the extra-cellular pathway, a tracer will only diffuse through the inter-endfoot gaps (pore space) and cannot enter the end-foot processes themselves (solid skeleton). Therefore, the diffusive transport across the endfoot sheath will be diminished by its gap area fraction, *ϕ*_g_. The endfoot processes are known to partially overlap [3], cf. Fig. 2 and the diffusive flux over the endfoot sheath is inversely proportional to the gap length (not endfoot sheath thickness), *l*_g_, estimated for capillaries at *l*_g_ = 0.45 µm [3]. The corresponding endfoot sheath thickness, *h*_ES_, is reported to be between 0.02 and 0.3 µm [3] in capillaries for chemically fixated tissue, while McCaslin et al. [5] report *h*_ES_ ≈ 1.0 µm for mouse cortex capillaries in-vivo and even larger *h*_ES_ for arterial and venous vessels. Since for geometrical reasons *l*_g_ ≤ *h*_ES_ (see Fig. 2), we propose *l*_g_ ≈ 1.5*h*_ES_ in the absence of quantitative in-vivo data. Due to obstructions in the endfoot gap channel (larger proteins, fibers, or gap junctions [14]) the effective diffusivity may be reduced by a factor *α*. Nicholson and Hrab ětová [22] demonstrate that in the extra-cellular space of the parenchyma where the inter-cellular space width is between 20-100 nm, the measured ratio of effective to free diffusivity is usually smaller than can be explained by the tortuosity of the pore space alone. For molecules with a hydrodynamic diameter that is one-tenth or more of the gap width, *α* needs to model size-dependent steric exclusion and restricted diffusion effects [23], for instance, with the Renkin equation [24, 25]. In particular, *α* and therefore diffusive permeability is zero for molecules much larger than the gap width.

The diffusive flux *F*_*D*_ [MT^−1^L^−2^] through the surface of a vessel segment can be computed as

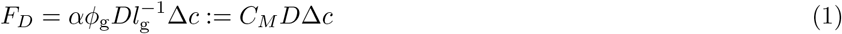

where Δ*c* is the concentration drop across the endfoot sheath, the gap area fraction *ϕ*_g_ is given by Eq. (5), *D* is the binary diffusion coefficient in aqueous solution, and

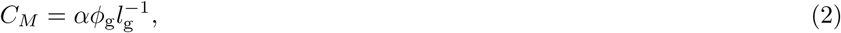

is the diffusion membrane coefficient (in units of m^−1^). Given a surface area *S* (for example for a tubular segment of length *l*_v_ and radius *r*_v_, *S* = 2*πr*_v_*l*_v_) with constant *C*_*M*_, we can compute the amount of a tracer crossing the endfoot sheath per unit time as *F*_*D*_*S*. The product *SC*_*M*_ *D* is sometimes called permeability-surface product or diffusion capacity [26, Ch. 10], in particular when referring to the surface integral of *C*_*M*_ *D* in a larger tissue portion. In this work, we normalize the diffusion capacity by the (free) binary diffusion coefficient *D*.

### Permeability for fluid flow

Although inter-endfoot gap slits are as narrow as *d*_g_ ≈ 20 nm, continuum theory of viscous flow can be applied for describing liquid water flow [27]. Furthermore, due to hydrophilic surface properties, we assume no-slip conditions at the cell membrane. We remark that the complex interface region (≈ 1 nm) between the endfoot’s lipid bilayer membrane and the bulk fluid adds some uncertainty to the effective width in addition to the uncertainty of width measurements and spatial variations. Therefore, we argue it is sufficient to approximate the hydraulic transmissibility by a simple parallel plate flow model as 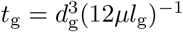, cf. [18], where *d*_g_ is the inter-cellular distance (i.e. gap width of the slit) and *µ* denotes the fluid viscosity. Using this expression, the flow rate *Q* [L^3^T^−1^] through the surface of a tubular segment of length *l*_v_ and radius *r*_v_ (and thus lateral surface area *S* = 2*πr*_v_*l*_v_) can be computed as

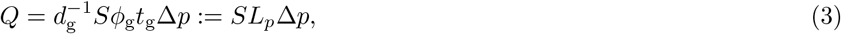

where Δ*p* is the effective pressure drop across the endfoot sheath (between extracellular space (ECS) and PVS) and the gap area fraction *ϕ*_g_ is given by Eq. (5) in terms of the vessel radius, *r*_v_, and

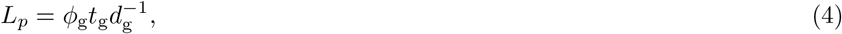

is the filtration coefficient (in units of m Pa^−1^ s^−1^). The product *SL*_*p*_ is also called filtration capacity [26, Ch. 11], in particular when referring to the surface integral of *L*_*p*_ in a larger tissue portion.

### Microvascular networks

We analyzed two microvascular networks (MVN) extracted from the mouse brain cortex in [19] as published in [28]. The raw vessel polylines extracted from segmented voxel images by [19] is smoothed by a Douglas-Peucker algorithm [29] using the local vessel radius as tolerance. For vessel classification (arteries and veins), blood pressure values (*p*) in every vessel segment are simulated with a finite volume method as described in [15] (but neglecting filtration across the blood-brain-barrier). The boundary conditions are based on estimations computed by Schmid et al. [30, 28]. We solve a modified Poiseuille-type flow using the in-vivo apparent viscosity relation proposed in [31] scaled to mouse red blood cells (using an average volume of 55 fL). The open-source software DuMu^x^ [32] was used as a finite volume solver with `dune-foamgrid` [33] for the network representation.

Using the computed pressure maps, vessel segments were classified as arterial vessels if its pressure exceeds the average pressure of all segments with *r*_v_ ≤ 4.5 µm (vessel radius excluding endfoot sheath), and as venous vessels otherwise. The networks and the obtained pressure distribution are shown in Appendix B (Fig. 9).

Since the network data is associated with vessel lumen radius data, *r*_v_, excluding the endfoot sheath and other perivascular structures but the tiling model is formulated in terms of the total outer radius (*r*_o_) of the astrocyte endfoot sheath, we require a model of how these radii are related. Based on data reported in [5], we assumed a thickness of the endfoot sheath, *h*_ES_, of 1 to 2.5 µm. Additionally, we chose the relation *h*_ES_ = 1 + 0.15(*r*_v_ − 3) modeling a linear increase with increasing vessel lumen radius. Moreover, larger vessels with *r*_v_ ≤3 µm are assumed to be sheathed by smooth muscle cells or ensheathing pericytes [34] located in between the endothelial layer and the endfoot sheath. Based on [34, Fig.3], we estimated the smooth muscle cell layer thickness to be approximately equal to *h*_ES_. This means ca. 1 µm for a pre-capillary arteriole with *r*_v_ = 3 µm and ca. 2 µm for a penetrating vessel with *r*_v_ = 10 µm. Finally, we added the thickness of the endothelial cell layer and basement membrane with 0.4 µm [35] for all vessels. In summary, *r*_o_ = 2*h*_ES_(*r*_v_)+0.4 = 1.3*r*_v_+1.5 for *r*_v_ ≥ 3 µm and *r*_o_ = *h*_ES_(*r*_v_)+0.4 = 1.15*r*_v_+0.95 otherwise. For the network analysis, the networks are split into 6 vertically stacked analysis layers (layer 0 being closest to the pial surface and layer 5 being closest to the white matter) of 200 µm thickness (100 µm for layer 5). Average values (*r*_v_, *r*_o_, *L*_*p*_, *C*_*M*_) have been computed as surface-area-weighted arithmetic averages of all vessels contained in the respective analysis layer.

## Results

### Astrocyte endfoot area distribution

Endfoot area distribution and resulting gap area fraction predicted by the model for 200 realizations with *r*_o_ = 2.9 µm^2^ (capillary) and *r*_o_ = 15.0 µm^2^ (venule and arteriole), with (corrected) mean endfoot area *A* shown in Fig. 3, are reported in Fig. 4. The resulting endfoot area distribution is well-modeled by a Gamma distribution^[5]^ (with mean 50 µm^2^ (capillary), 110 µm^2^ (venule) and 490 µm^2^ (arteriole), respectively). The resulting gap area fraction distribution is well-approximated by a normal distribution with mean gap area fractions of 0.0056 (capillary), 0.0038 (venule), and 0.0018 (arteriole). For comparison with [3, 18], we also report the resulting number of gaps counted per vessel cross-section (‘gaps/ring’) on average, which is lowest in the capillary (3.2), highest in venules (11.5), and intermediate in arterioles (5.4).

**Figure 4.**
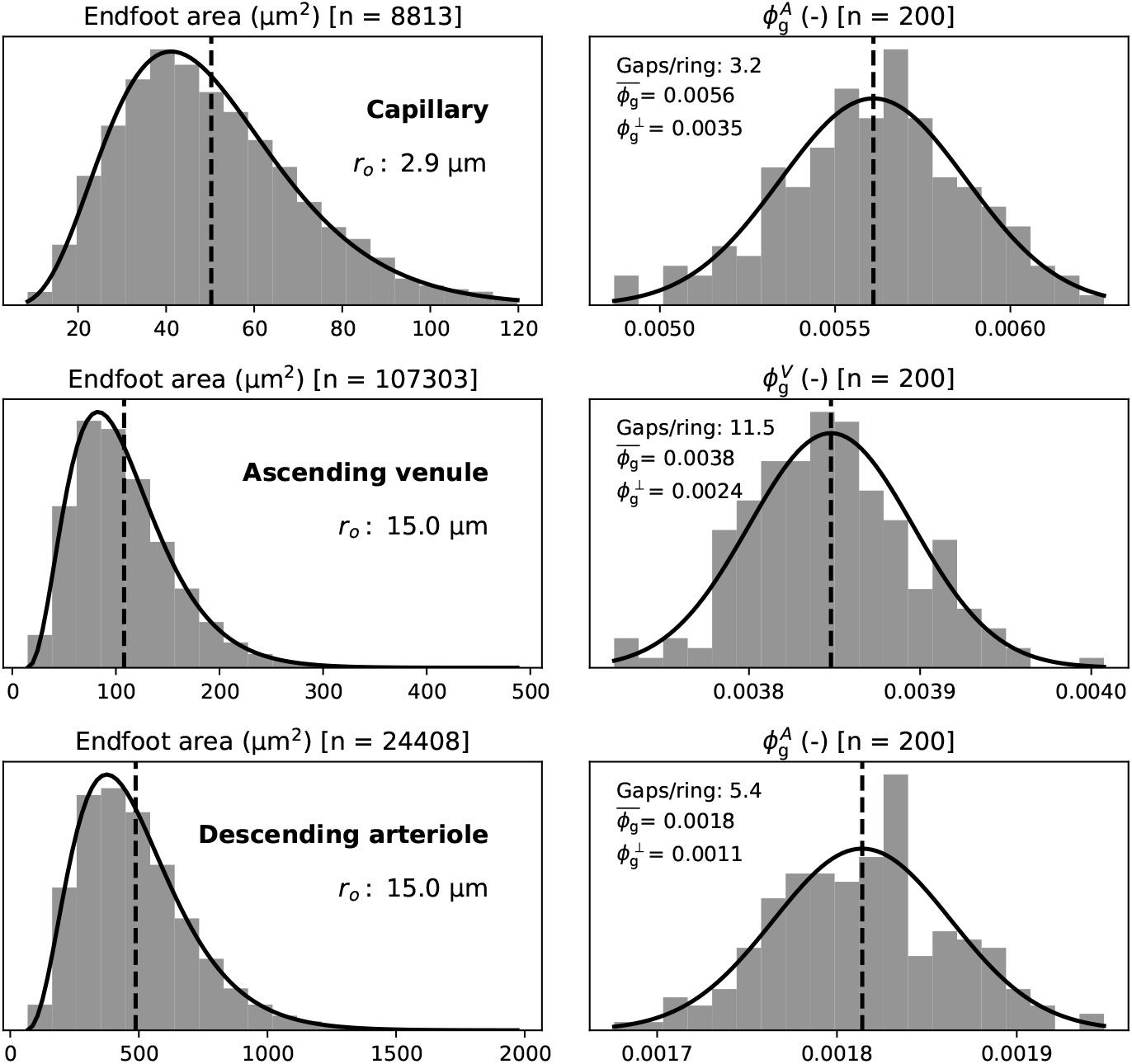
Endfoot area and gap area fraction from Voronoi tessellations. (First row) endfoot area distribution (left) and resulting gap area fraction (right) for 200 realizations with *r*_o_ = 2.9 µm^2^ (capillary), (second row) *r*_o_ = 15.0 µm^2^ (venule), (third row) *r*_o_ = 15.0 µm^2^ (arteriole). The dashed vertical line marks the mean (capillary: *A* = 50 µm^2^, 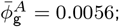 venule: *A* = 110 µm^2^, 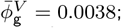 arteriole: *A* = 490 µm^2^, 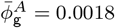). The quantity ‘gaps/ring’ states the number of inter-cellular endfoot gaps, on average, on cross-sectional vessel cuts. The resulting gap area fraction—if this value were to be extrapolated to the total surface—is denoted by 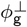. ‘Gaps/ring’ and 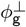 are reported for comparison with experimental data. The solid lines represent fitted continuous distributions using a Gamma distribution for the endfoot area and a normal distribution for the gap area fraction.

Additionally, the model was run for 50 different diameters with model-corrected *A* (shown in Fig. 3). For each diameter, we generated 20 samples (a total of *n* = 1000 samples). The resulting data including mean, and 5th, 25th, 75th, and 95th percentile are reported in Fig. 5 for both venous and arterial vessels.

**Figure 5.**
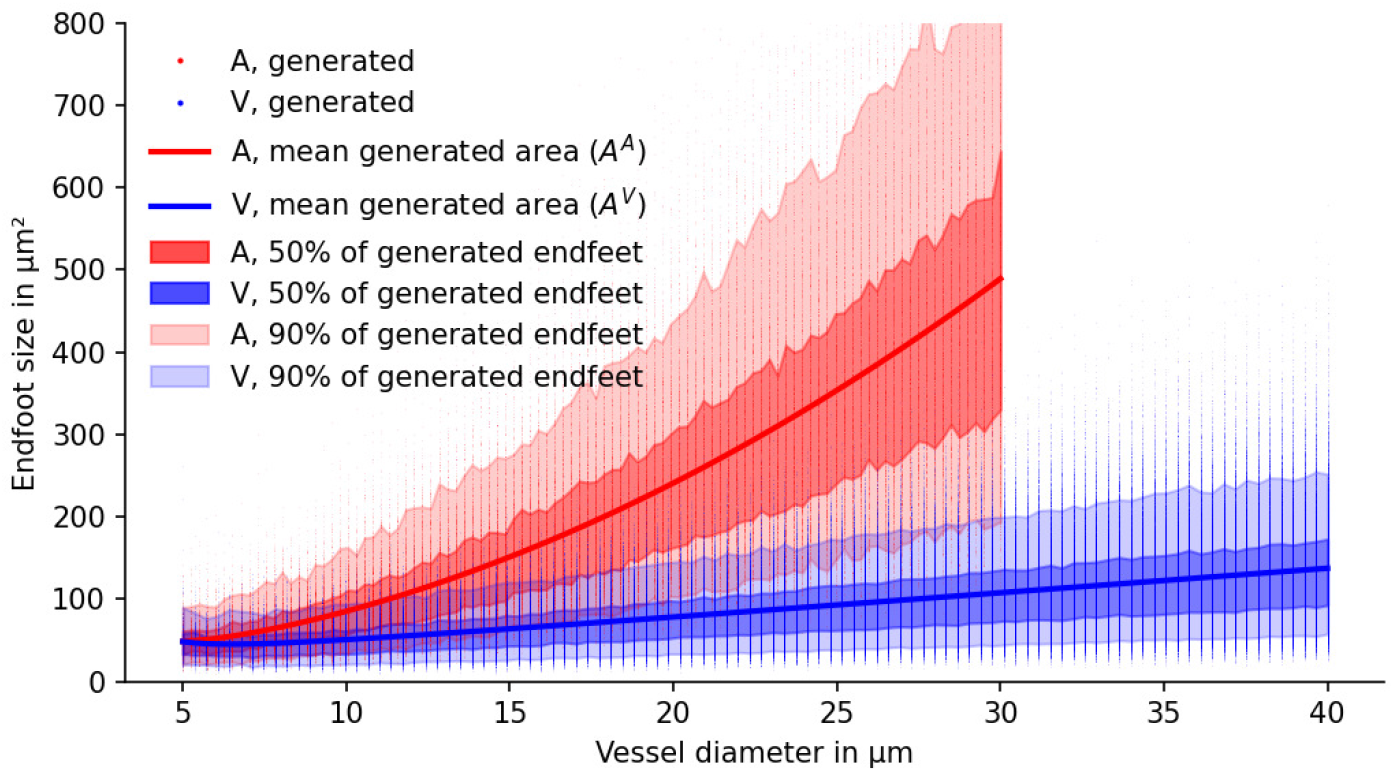
Endfoot areas of generated Voronoi-diagram-based endfoot patterns. Given the mean endfoot areas estimated in Fig. 3 (solid lines), *n* = 1000 realizations were generated for varying vessel diameters. Individual dots correspond to a single astrocyte endfoot of a realization. The shown mean corresponds to *A*^*A*^ and *A*_*V*_. Individual endfoot coverage areas show large variability (in agreement with what is reported in [6, Fig.2d]) and follow a Gamma distribution for a given vessel diameter (and mean endfoot area), cf. Fig. 4.

For comparison with previously published data, we computed based on the diameter-area relations that on average, small vessels (*r*_*v*_ < 4.5 µm, average not weighted by radius prevalence in a network) have endfoot density of ca. 1.9 × 10^4^ endfeet per mm^2^ surface area. Larger venous vessels (*r*_*v*_ > 4.5 µm) show ca. 1.0×10^4^ endfeet/mm^2^ and larger arterial vessels (*r*_*v*_ > 4.5 µm) show the lowest density of ca. 0.4 × 10^4^ endfeet/mm^2^.

### Gap area fraction for different vessels

Using the same *n* = 1000 samples as for the data in Fig. 5, in combination with the gap width and length reported by Mathiisen et al. [3], we computed the resulting gap area fraction *ϕ*_g_ for each realization. The results are shown in Fig. 6. For small diameters (capillaries), the gap area fraction for venous vessels, 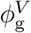, and arterial vessels, 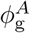, are similar, while for increasing vessel diameters 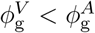.

**Figure 6.**
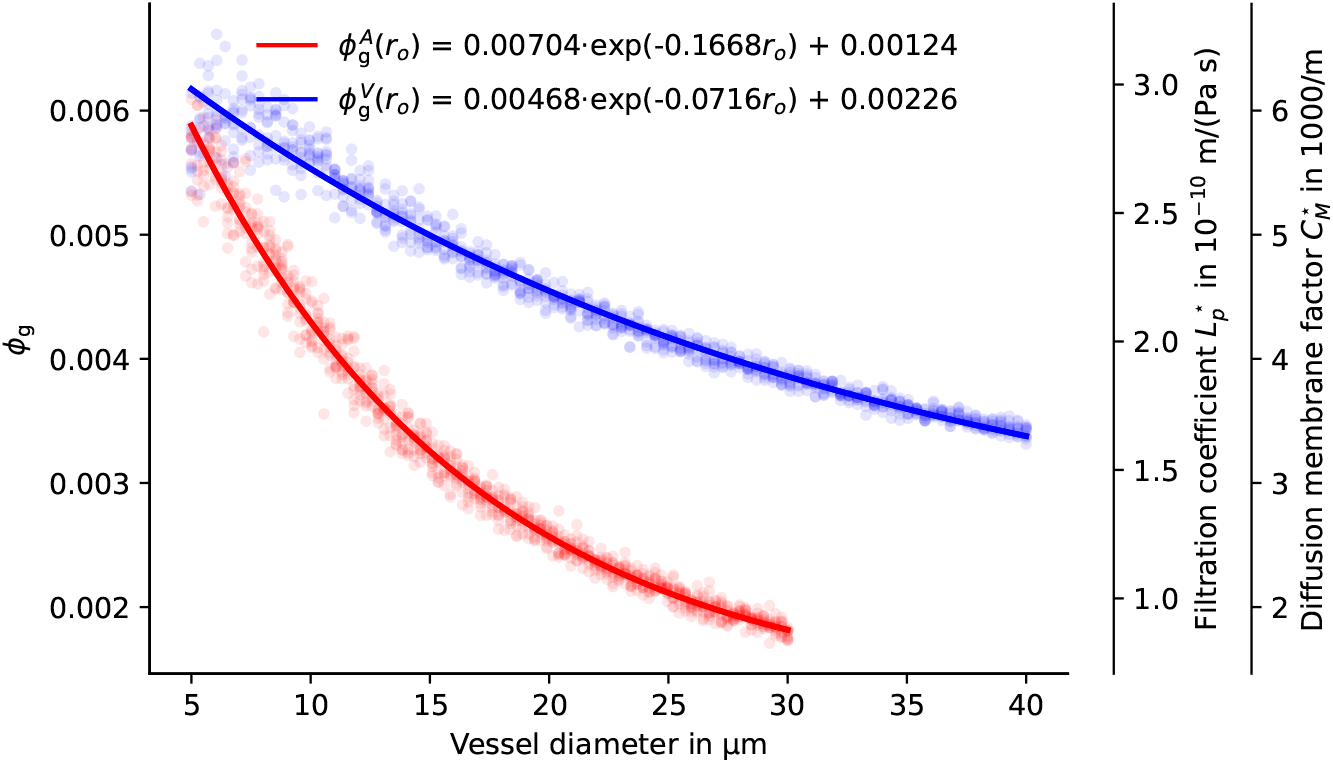
Functional relation between vessel diameter and gap area fraction. Gap size fraction from generated endfoot sheath patterns for uniform gap width of *d* = 20 nm. Dots show individual realizations (*n* = 800 for each arterial and venous vessel). Solid lines are exponential curve fits (*r*_o_ = 0.5*d*_o_ is the vessel radius including the endfoot sheath). Estimates for 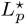 and 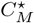 (right axes) are computed with *l*_g_ = 1 µm, a fluid viscosity of *µ* = 0.69 × 10^−3^ Pa s and *α* = 1. Both 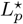 and 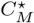 are proportional to *ϕ*_g_*/l*_g_.

In summary, the model predicts 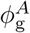 and 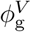 given mean endfoot area, *A*, vessel radius including endfoot sheath, *r*_o_, and gap with, *d*_g_. Using a gap width of *d*_g_ = 20 nm [14, 3], we obtain the empirical relations,

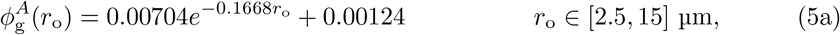

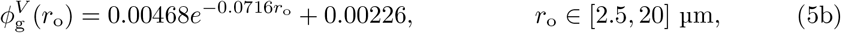

where the radius *r*_o_ is inserted in µm, cf. Fig. 6.

Based on a constant gap length, *l*_*g*_, the gap area fraction for the smallest capillary vessels is about 3 times higher than that of 15 µm radius arterial vessel, and about 2 times higher than that of venous vessel of the same caliber. The smaller increase in endfoot size reported by [6] for venous vessels in comparison with arterial vessels leads to effectively higher gap area fractions in venous vessels with increasing vessel radius.

### Permeability for diffusive transport of passive tracers

The resulting estimates for *C*_*M*_ using an obstruction of *α* = 1 (i.e. no obstructions; smaller *α* values would decrease the *C*_*M*_ estimates) and constant *l*_g_ = 1 µm are shown in Fig. 6. Also taking into consideration the variation of *h*_ES_ in the microvascular networks (see Methods), estimates range between *C*_*M*_ ≈ 500 m^−1^ for the largest arterioles and *C*_*M*_ ≈ 6000 m^−1^ for the smallest capillaries.

### Permeability for fluid flow

The resulting estimates for *L*_*p*_ using a viscosity of *µ* = 0.7 × 10^−3^ Pa s (water at 37 °C, larger assumed viscosity values would increase the *L*_*p*_ estimates) and constant *l*_g_ = 1 µm are given in Fig. 6. Also taking into consideration the variation of *h*_ES_ in the microvascular networks (see Methods), estimates range between *L*_*p*_ ≈ 0.2 × 10^−10^ m Pa^−1^ s^−1^ for the largest arterioles and *L*_*p*_ ≈ 3 × 10^−10^ m Pa^−1^ s^−1^ for the smallest capillaries, cf. Fig. 7.

**Figure 7.**
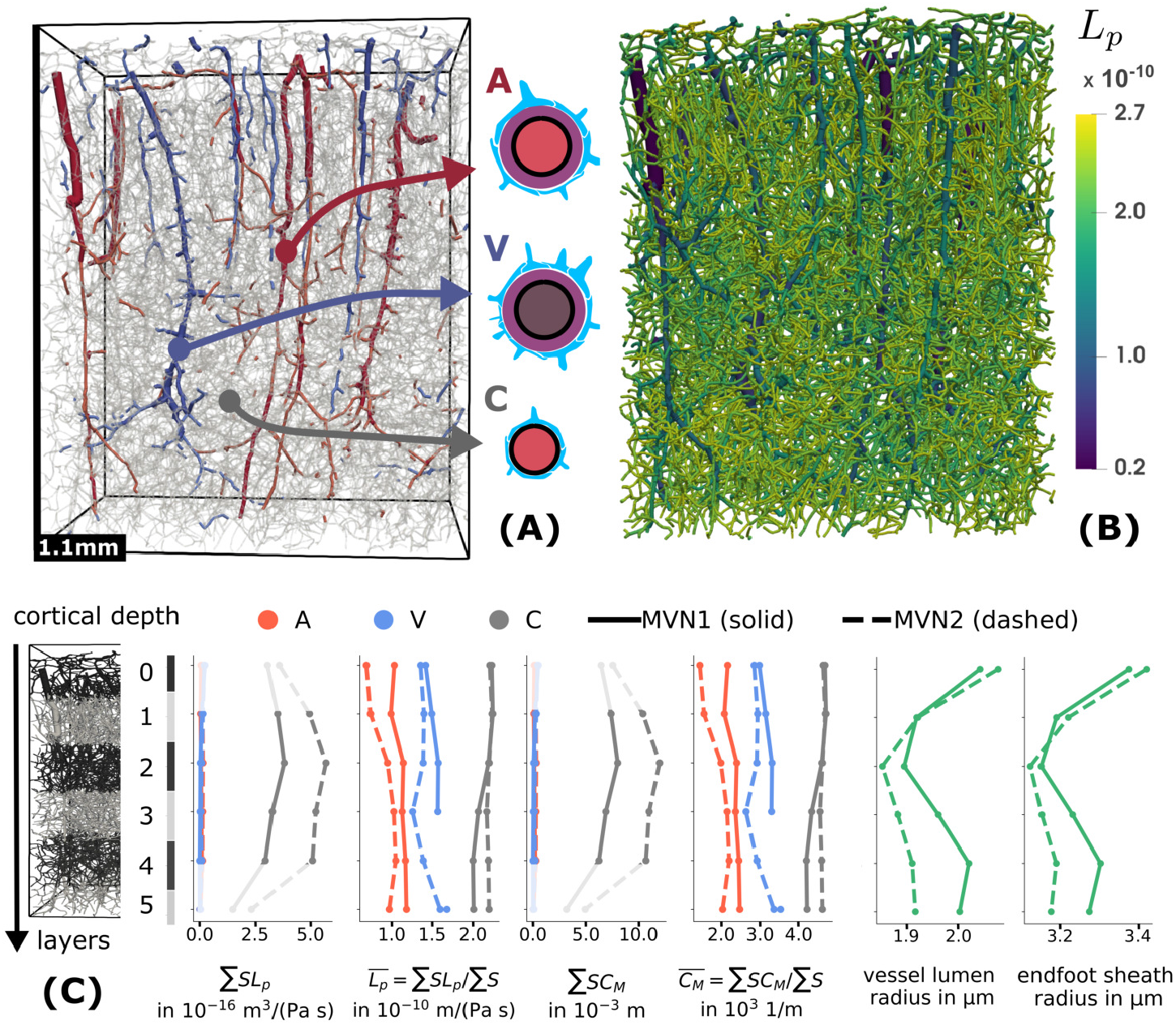
Microvascular networks (MVN). MVN1 and MVN2 are networks extracted from mouse cortex data [19]. The vessel segment visualizations show MVN2. Pial vessels have been removed. Classification into arterial and venous vessels is based on pressure maps computed with a vascular graph model (see Methods and Fig. 9). (A) Vessels with *r*_v_ ≥ 3 µm are shown in red (arterial) and blue (venous); smaller vessels are translucent gray. Segments are rendered as cylinders with radius *r*_v_. (B) The color map is scaled by the estimated filtration coefficients *L*_*p*_ shown for each vessel segment. Segments are rendered as cylinders with radius *r*_o_. (C) For each of the 6 analysis layers (layer 0 is closest to the pial surface) of 200 µm vertical thickness (100 µm for layer 5) and both networks, the filtration capacity, the average *L*_*p*_, the diffusion capacity, and the average *C*_*M*_ for each vessel category (A: arterial, V: venous, C: capillary vessels) is shown. The surface area of each vessel segment has been computed by assuming a cylinder with radius *r*_o_. The averaged segment lumen radius (*r*_v_ from [28]) and the averaged estimated outer endfoot sfheath radius (*r*_o_) over depth are shown in the rightmost figure. Missing data points correspond to ∑*S* = 0.

### Microvascular networks

We analyzed two microvascular networks from the mouse brain cortex [19], labeled MVN1 and MVN2. Volumes and surfaces computed by assuming cylindrical segments with estimated outer radius *r*_o_ (including endfoot sheath) and vessel lumen radius *r*_v_ (from [28]) are given in Table 1. MVN2 has a 18 % larger vessel volume fraction (2.8 % and 3.4 % including endfoot sheath) and an 10 % larger surface-to-volume ratio than MVN1 (1.7×10^4^ m^2^ m^−3^ and 1.9×10^4^ m^2^ m^−3^). In both networks, the surface area of small vessels (*r*_*v*_ < 3 µm) exceeds the area of the larger vessels by a factor 10 or more.

**Table 1.** Average parameters computed for two microvascular networks. MVN1 and MVN2 are networks extracted from the mouse cortex [19]. Capillaries (C) are defined as vessels with *r*_v_ < 3.0 µm. Venous (V) and arterial (A) vessel segments (*r*_v_ ≥ 3.0 µm) are distinguished by local blood pressure (see Methods). Pial vessels are excluded from the analysis. Outer surface refers to the endfoot sheath surface. The surface area is computed on the basis of circular cross-secitons and estimated *r*_o_ (outer radius including endfoot sheath, see Methods). Surfaces and volumes are based on the assumption of cylindrical segments with radius *r*_v_ (lumen, *L*) or *r*_o_.

| Symbol | MVN1 | MVN2 | unit | description |
| --- | --- | --- | --- | --- |
| $S_C$ | $8.5 \times 10^{-6}$ | $1.2 \times 10^{-5}$ | $\text{m}^2$ | endfoot sheath surface area (C) |
| $S_A$ | $3.2 \times 10^{-7}$ | $7.5 \times 10^{-7}$ | $\text{m}^2$ | endfoot sheath surface area (A) |
| $S_V$ | $2.6 \times 10^{-7}$ | $4.6 \times 10^{-7}$ | $\text{m}^2$ | endfoot sheath surface area (V) |
| $S$ | $9.1 \times 10^{-6}$ | $1.3 \times 10^{-5}$ | $\text{m}^2$ | endfoot sheath surface area (all) |
| $V$ | $5.5 \times 10^{-10}$ | $6.9 \times 10^{-10}$ | $\text{m}^3$ | total sample volume (bounding box) |
| $\zeta$ | $2.8 \times 10^{-2}$ | $3.4 \times 10^{-2}$ | - | volume fraction vessel (outer) |
| $\zeta^L$ | $1.0 \times 10^{-2}$ | $1.2 \times 10^{-2}$ | - | volume fraction vessel (lumen) |
| $S/V$ | $1.7 \times 10^4$ | $1.9 \times 10^4$ | $\text{m}^2 \text{ m}^{-3}$ | surface to volume ratio (outer surface) |
| $S^L/V$ | $1.0 \times 10^4$ | $1.2 \times 10^4$ | $\text{m}^2 \text{ m}^{-3}$ | surface to volume ratio (lumen surface) |

For each vertical depth analysis layer (see Methods), the filtration and diffusion capacity as well as averaged filtration and diffusion membrane coefficients are shown in Fig. 7. While the filtration coefficient *L*_*p*_ (and similarly the diffusion membrane coefficient, *C*_*M*_) in individual segments differs by a factor 10 between the largest arteriole segments and the smallest capillaries, the layer-averaged coefficient only varies by a factor 2. The surface-weighted averages over the entire network are found to be *L*_*p*_ = 2.1×10^−10^ m Pa^−1^ s^−1^ and *C*_*M*_ = 4.4×10^3^ m^−1^, with only 1 % difference between MVN1 and MVN2. The total filtration capacity per tissue volume is found to be 3.4 × 10^−6^ m Pa^−1^ s^−1^ m^−1^ (MVN1), 4.1× 10^−6^ m Pa^−1^ s^−1^ m^−1^ (MVN2). The total diffusion capacity per tissue volume is found to be 7.2 × 10^7^ m^−2^ (MVN1), 8.5 × 10^7^ m^−2^ (MVN2). The filtration and diffusion capacity peaks in layer 2, where it is between (10 to 14 % larger than in layer 1, 3, and 4 (0 and 5 are excluded from this comparison due to differences in layer size and occupied volume). This coincides with the lowest average vessel radius, cf. Fig. 7.

## Discussion

### Endfoot gap area fraction

In [3], the average number of endfoot gaps per capillary cross-section is reported as 2.5 (2.3 to 2.9 in 3 different animals). For the modeled capillary with *r*_o_ = 2.9 µm, the predicted number (‘gaps/ring’ in Fig. 4) of 3.2 is only slightly larger (20 %). However, without correction of the projection error (Appendix A), capillary endfoot size is estimated in [6] at only 25 µm^2^ corresponding to about 4.0 ‘gaps/ring’ (simulated with our model). Hence, the correction by the model allows us to partially resolve an apparent mismatch between the data reported by [3] and [6]. The comparison may be further affected by the different measurement methods employed by [3] and [6], measurement errors, and the quality of the area correction computed by our model. Finally, there might be inter-species variations between rats and mice.

By extrapolating ‘gaps/ring’ and the gap width of *d* = 20 nm to all of the surface, the authors of [3] conclude that about 0.3 % of the endfoot sheath surface is comprised of gaps—a number also used by [18] to estimate endfoot sheath permeability. We note that this computation effectively assumes that gaps run parallel to the longitudinal vessel axis. Under this assumption, we compute for capillaries, a reduced gap area fraction 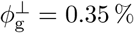 matching well the value obtained in [3]. However, the corresponding actual gap area fraction estimated by our surface tessellation model is *ϕ*_g_ = 0.56 %, cf. Fig. 4, and therefore significantly larger. The latter value can be considered an improved estimate correcting for yet another projection error.

For a simple estimate of gap area fraction, we may assume a regular hexagonal tiling for *A* = 50 µm^2^ and *d* = 20 nm corresponds to a value *ϕ*_g_ = 0.0052^[6]^ (to be compared with 0.0056) for the capillary and *ϕ*_g_ = 0.0035 (compared with 0.0038) for the vein of the same caliber analyzed in Fig. 4. However, regular tiling falls short of providing a model for individual endfoot size variability.

With respect to the variation with vessel type, we remark that assuming constant gap width for all vessels results in a linear correlation between filtration and diffusion membrane coefficient (both quantities depend linearly on the gap area fraction). Therefore, differences in permeability result from variations in the gap area fraction rather than individual gap anatomy. Such a correlation is, for example, also observed for the endothelium of different capillary types [26, Ch. 10.6].

However, there is significant uncertainty regarding both gap width in-vivo and general astrocyte coverage. The estimates in this work consider a continuous coverage with astrocyte endfeet of all microvessels [5]. Firstly, the actual coverage may be reduced with, for example, astrocyte bodies or microglia substituting endfoot processes on the vessel surface. If the inter-cellular gap size is not significantly altered, the provided estimates by our model still hold. Secondly, both Mathiisen et al. [3] and Wang et al. [6] worked with chemically fixated tissue. Korogod et al. [39] compared cryogenic and chemical fixation techniques, and report significant differences in the resulting endfeet cavity fraction (37 % vs. 4 %). At cavity fractions this large, the astrocyte endfoot sheath would be irrelevant in terms of a proposed barrier function. This result is contrasted by in-vivo observation of continuous coverage [2, 5, 40]. Additionally, Kubotera et al. [41] observed that after laser ablation astrocytes restore the endfoot coverage of microvessels in-vivo. (Mills and coworkers call this tendency to re-cover blood vessels after disruptions *endfoot plasticity* [42].) On the other end, the effective gap area fraction is reduced, if inter-endfoot gap junctions (2-3 nm [14]) are found to be present in-vivo with significant density (neglected in this work). It is also reduced for molecules whose hydrodynamic radius is a significant fraction of the gap width (modeled by the parameter *α*).

Apart from structural uncertainty, astrocytes are known to change their volume under varying conditions [43]. Changes in cell sizes and changes in the radius of the endfoot sheath could alter its hydrodynamic properties—a potential regulatory mechanism of fluid flow and substance transport [44, 6]. We are not aware of quantitative data describing how the endfoot gap width *d*_g_ or endfoot sheath thickness *h*_ES_ changes with such alterations. If such data became available, Eqs. (2) and (4) allow estimating the effect of alterations on the gap area fraction (and *C*_*M*_, *L*_*p*_). Moreover, vessel diameters are highly dynamic and can dilate up to 30 to 40 % of the vessel diameter [45] which leads to mechanical deformation of the astrocyte endfoot sheath observed in-vivo [46].

To the best of our knowledge, direct evidence for full coverage (or its absence), a precise inter-endfoot gap width quantification in-vivo and its variation, as well as quantitative data on temporal dynamics are still lacking. The fluid flow rate 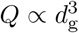 and the diffusive flux *F*_*D*_ ∝ *d*_g_ strongly depend on the assumed gap width *d*_g_ resulting in a large (maybe the largest) source of uncertainty for the estimation of extra-cellular transport across the endfoot sheath.

### Permeability for diffusive transport of passive tracers

The estimated permeability of the endfoot sheath can be compared to adjacent barriers and tissues. The vessel wall is assumed to be virtually impermeable to many molecules. In diseased tissue, for example, neurodegenerative diseases such as multiple sclerosis (MS) or glioma higher permeability has been observed in lesion tissue. For example, MRI contrast agents such as gadobutrol (*D* ≈ 3.5 × 10^−10^ m^2^ s^−1^ [15]) can leak out of blood vessels in MS lesions or glioma tissue. In [15], *C*_*M*_ *D* ≈ 1 × 10^−7^ m s^−1^ has been estimated for Gadobutrol leakage across the vessel wall in MS lesions, corresponding to *C*_*M*_ = 300 m^−1^ which is an order of magnitude smaller than average values obtained in this work for the endfoot sheath, cf. Fig. 7. For skeletal muscle microvascular walls and small hydrophilic molecules, *C*_*M*_ has been estimated at *C*_*M*_ ≈ 100 to 200 m^−1^ [23, Fig.2]. The brain cortex microvascular vessel walls are commonly assumed to be orders of magnitude less permeable than in skeletal muscle. This signifies that the endothelial layer is a much less permeable barrier than the astrocyte endfoot sheath, where we estimated the lowest *C*_*M*_ for large penetrating arterioles with *C*_*M*_ ≤ 1000 m^−1^ and values up to *C*_*M*_ ≤ 6000 m^−1^ for capillaries.

The estimated permeability can also be compared to that of the extra-cellular space (ECS). To this end, we consider a 1 µm thick slab of ECS. With a porosity of 0.2 and tortuosity factor of 0.35 [47, 1*/λ*^2^], we obtain *C*_*M*_ = 7 × 10^4^ m^−1^. Hence, the endfoot sheath is more than an order of magnitude less permeable than the ECS given a slab of comparable thickness. Therefore, the endfoot sheath could locally act as a barrier. It could also promote the compartmentalization of substances, depending on whether low-permeability perivascular pathways parallel to the vessel exist in the vicinity.

We remark that lower permeability does not necessarily mean slower transport. The magnitude of diffusive transport depends on the concentration drop Δ*c* as well, cf. Eq. (1). As vessel structures constitute thin tubular sources (in an infiltration scenario) or sinks (in a clearance scenario), the magnitude of the concentration gradient is much larger in the vicinity of the vessels and quickly decays with distance. Therefore, the question as to whether the effect of a lower local permeability is significant for a given scenario goes beyond the scope of the present work.

### Permeability for fluid flow

In the brain cortex microvasculature, the filtration coefficient of the vessel wall is thought to be very low. Kimura and coworker [48, Tab.3] measured *L*_*p*_ = 2.8 to 4.1× 10^−12^ m Pa^−1^ s^−1^ in single rat brain arterioles. Fraser and Dallas [49] report *L*_*p*_ = 2 × 10^−13^ m Pa^−1^ s^−1^ in frog brain microvessels. A 1 µm slab of ECS corresponds to a *L*_*p*_ value of approximately 0.5 to 3×10^−8^ m Pa^−1^ s^−1^ [50, 51, 52]^[7]^ or larger^[8]^. We estimated the lowest *L*_*p*_ values for large arterioles with *L*_*p*_ ≈ 2.0×10^−11^ m Pa^−1^ s^−1^, and the largest values for capillaries, *L*_*p*_ ≈ 2.0 × 10^−10^ m Pa^−1^ s^−1^.

Hence, similar to the results for diffusion, the endfoot sheath filtration coefficient is one order of magnitude larger than that of the vessel wall. On the other hand, it is two orders of magnitude smaller than a slab of ECS of similar thickness, making the astrocyte endfoot sheath a limiting component for the extra-cellular PVS-ECS exchange of fluids.

Using the same parallel plate model as for *L*_*p*_, Eq. (4), we can estimate Péclet numbers for transport through the gaps as 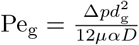 (using *l*_g_ as characteristic length). The Péclet number estimates the importance of advective transport over diffusive transport across the end foot sheath. Given the difference in permeability to that of the vessel wall, across which pressure drops on the order of 1000 Pa may occur due to filtration across the capillary endothelium (estimated for example with the models presented in [55, 15]), we assume a pressure drop (Δ*p*) across the endfoot sheath of 10 Pa to 100 Pa. Since typical binary diffusion coefficients in aqueous solution are in the order of 1 × 10^−9^ m^2^ s^−1^, transport of solutes across the endfoot sheath is dominated by diffusion (Pe_g_ ≈ 0.003 ≪ 1).

In [9], Asgari and coworkers estimate an inter-endfoot-gap permeability of *L*_*p*_ = 1.35 × 10^−10^ m Pa^−1^ s^−1^ (based on a capillary circumference 15.7 µm, endfoot thickness *h* = 1 µm and a parallel plate model, and conversion based an endfoot area of 78 µm^2^). This matches well with the average value estimated in arterioles in this work but is lower by about half what we estimate for capillaries. The difference can be explained by the assumption of Asgari that the assumed representative endfoot fully wraps around the vessel (‘gaps/ring’ is 1) which results in a lower gap area fraction.

In [18], Vinje and colleagues estimate comparable parameters for human brain tissue. In particular, the authors estimated the endfoot sheath resistance (inverse permeability) around arterial and venous vessels (excluding capillaries). The reported resistances correspond to *L*_*p*_ = 2 × 10^−10^ m Pa^−1^ s^−1^ for arterial and *L*_*p*_ = 3 × 10^−10^ m Pa^−1^ s^−1^ for venous vessels. The numbers are, in part, based on the gap area fraction estimate provided in [3] for capillaries in rats. As explained above, this number (based on the quantity ‘gaps/ring’) results in an underestimation of *ϕ*_g_ and therefore *L*_*p*_ of ca. 35 % and the suggested higher values would be *L*_*p*_ ≈ 3 × 10^−10^ m Pa^−1^ s^−1^ for arterial and *L*_*p*_ = 4.5 × 10^−10^ m Pa^−1^ s^−1^ for venous vessels. However, we used the endfoot thickness distribution estimated by [5] based on in-vivo mouse brain data, whereas a constant size straight channel model with *l*_*g*_ = *h*_ES_ = 1 µm is used by [18]. Thus, our resulting permeability for arterioles and venules are approximately half the values of [18], respectively, cf. Fig. 7.

### Microvascular networks

For the two considered microvascular networks, we find that the filtration and the diffusion capacity are largest at about 40 % of cortical depth. This layer also shows the smallest average vessel diameters, cf. Fig. 7, and a significant peak in neuron density [56]. A high endfoot density per surface area as in the capillaries, cf. [5], corresponds to a higher permeability of the endfoot sheath due to an increase in the gap area fraction. The average filtration and membrane diffusion coefficients are dominated by the average values for capillaries and appear to be independent of depth. Hence, the increased filtration capacity at 40 % seems to be a result of an increased surface area rather than an increased vessel permeability. This matches with the observation that vessel density is largest in this cortical layer [19, 45, 57].

To the best of our knowledge, the variability of endfoot sizes in the endfoot sheath has not been analyzed using microvascular networks comprising all vessels in a given tissue portion before. Based on the distribution of penetrating arterioles and venous from the macaque cortex [58], Vinje et al. [18] estimate the surface permeability product of the human brain (using an approximate human brain volume of *V* ≈ 1 L). If normalized by the sample volume to eliminate the effect of spatial scale, their estimate corresponds to the volume-specific quantities 2.2 × 10^−7^ Pa^−1^ s^−1^ for arterioles and 2.0 × 10^−7^ Pa^−1^ s^−1[9]^ for venules, while we obtain 6.6 × 10^−8^ Pa^−1^ s^−1^ (MVN1), 6.6 × 10^−8^ Pa^−1^ s^−1^ (MVN2) for arterioles, and 9.5 × 10^−8^ Pa^−1^ s^−1^ (MVN1), 9.1 × 10^−8^ Pa^−1^ s^−1^ (MVN2) for venules. The difference is expected since we estimated lower endfoot sheath permeability.

Although not directly significant for the permeability of the endfoot sheath (but relevant for propositions about its main function), we additionally provide cell density estimates resulting from the analysis of the microvascular networks in combination with astrocyte endfoot areas. The assumed diameter-area relations mean that on average, small vessels (*r*_*v*_ < 3.0 µm) show an average endfoot density of 2 × 10^4^ endfeet/mm^2^ surface area. Larger venous vessels (*r*_*v*_ > 3.0 µm) show 1 × 10^4^ endfeet/mm^2^ and larger arterial vessels (*r*_*v*_ > 3.0 µm) show the lowest density of 0.4 × 10^4^ endfeet/mm^2^. McCaslin and coworkers [5] find 1 × 10^4^ endfeet/mm^2^ for capillaries, 0.4 × 10^4^ endfeet/mm^2^ for venules, 0.3 × 10^4^ endfeet/mm^2^ for arterioles in-vivo in mouse cortex.

Using the endfoot sheath surface areas in Table 1 and our density estimates, we compute about 170 000 (MVN1), 245 000 (MVN2) endfeet around small vessels, 2600 (MVN1), 4600 (MVN2) endfeet around larger venous vessels, and 1300 (MVN1), 3000 (MVN2) endfeet around larger arterial vessels. 97 % of endfoot processes are therefore expected to be around capillaries. Using an estimate of astrocyte densities in the mouse cortex (20000 ± 13000 cells/mm^3^ [56]) this means the domain of the analyzed networks contains about 11 000±7000 (MVN1) and 14 000±9000 (MVN2) astrocytes with 16 (MVN1) and 19 (MVN2) endfoot processes per astrocyte on average.

Finally, we want to stress that with regard to the prediction of transport across or in parallel to the endfoot sheath, in addition to the presented permeability parameters, a dynamic model for pressure and concentration around vessel networks on the µm to mm scale (meso-scale) is needed. Concerning implication for macroscale transport models (organ-scale), we remark that the integral values reported here for the microvascular networks may be used as a starting point to estimate parameters for tissue transport models based on homogenization or mixture theory. However, one should be aware that effective filtration and diffusion capacity on the macro-scale generally depend on the local meso-scale pressure and concentration distributions which is an unresolved issue of such models [55] in the context of tissue perfusion simulations.

### Relevance in the light of the glymphatic theory

Cerebrospinal fluid (CSF) flow through perivascular spaces is a crucial component of the recently proposed glymphatic theory [59]. Pial perivascular CSF flow has been observed and quantified in [60]. Furthermore, intake of various tracers (Dextran, Gadobutrol) into the parenchyma has been reported to be modified by sleep and disease in both mice and humans [61, 62, 63]. Crucial to determining the mechanisms involved in the intake is to determine the type and magnitude of fluid flow and molecular transport along the different pathways: perivascular, intra-cellular, and extra-cellular; and the resistance of barriers between these compartments and the resistance of efflux pathways. Therefore, the herein presented permeability estimates for the astrocyte endfoot sheath being a component of all conceived pathways, provide a starting point for estimating diffusive and advective fluxes outside of the microvasculature.

In [6], the authors estimate the effect of varying astrocyte endfoot gap density on transmembrane CSF flux based on (at least) three assumptions^[10]^: (1) there is a fluid-filled connected perivascular space (PVS) from descending arterioles all the way down to capillaries; (2) there is a net CSF flow within the PVS from the cortical surface into the capillary bed driven by axial pressure gradients in the PVS; (3) water transport across the endfoot sheath (or transport through intra-cellular pathways) does not affect the pressure distribution in the PVS, i.e. the exchange is small in comparison to the perivascular flow rates. In a theoretical analysis based on these assumptions, the authors conclude that varying endfoot gap fractions help “maintaining perivascular-interstitial flux through the cortical depth” [6]. This conclusion is based on the comparison of a single arteriole with a single capillary within a hierarchical network: the arteriole has a lower surface-specific permeability (*L*_*p*_) but experiences a larger pressure drop (Δ*p*) across the endfoot sheath than the capillary (given the authors’ assumptions). These competing effects cancel each other out, so that the resulting local fluxes across the endfoot sheath are approximately equal in both vessels. However, we want to additionally point out that for a given portion of tissue (as in Fig. 7), since there is many more capillaries than arterioles with a much higher total surface area (Table 1), perivascular-interstitial exchange (even with the authors’ assumptions) would happen predominantly around capillaries^[11]^. Moreover, the latter statement remains true, even if the arteriole endfoot sheath would have the same (higher) *L*_*p*_ as the capillaries. However, regardless of this remark, the low permeability (high resistance) of the endfoot sheath in comparison with the ECS may lead to slightly enhanced fluid flow parallel to vessels within the PVS (under the premise that a sufficient driving force and a connected pathway exist).

## Conclusion

This work shows how a data-informed theoretical model of astrocyte endfoot size distributions (based on Voronoi tessellations) can be used to relate data from various experimental and theoretical works and arrive at estimates for the endfoot sheath permeability and its variation in microvascular networks from mouse brain cortex. We estimated filtration coefficients in the range *L*_*p*_ = 0.2×10^−10^ m Pa^−1^ s^−1^ to 2.7× 10^−10^ m Pa^−1^ s^−1^ (average 2.1 × 10^−10^ m Pa^−1^ s^−1^) and diffusion membrane coefficients in the range *C*_*M*_ = 0.5 × 10^3^ m^−1^ to 6 × 10^3^ m^−1^ (average 4.4 × 10^3^ m^−1^). This means that the astrocyte endfoot sheath is more than one order of magnitude more permeable than the vessel wall but about two orders of magnitude less permeable than a similarly thick layer of extra-cellular space. The numbers are complemented by formulas such that they can be adapted in the case that other data becomes available. In particular, we estimated a relation between the inter-endfoot gap area fraction and the vessel radius given a constant gap width and find values in the range of 0.2 % to 0.6 %. The data is presented with the intent to be useful for detailed modeling studies of transport of substances in the brain cortex including microvascular network architecture. The estimates are based on the assumption of continuous endfoot coverage of cortical micro-vessels in mice with an approximately constant inter-endfoot gap width of 20 nm and largest uncertainty for the permeability of the extra-cellular pathway stems from the absence of direct evidence of continuous endfoot coverage and the precise geometry of inter-endfoot gaps in-vivo.

## A Underestimation of endfoot area by 2D image analysis

The analysis of varying endfoot area on the surface of vessels of different caliber by [6] is based on two-dimensional image analysis. A (2D) image of a vessel is taken. This image looks like Fig. 1 (bottom), cf. [6, Fig.2]. Based on the image inter-endfoot gaps are segmented and the area surrounded by inter-endfoot gaps is identified as endfoot area. There is two main error inherent to the methodology:

1. (Projection error) The image shows the (originally curved) vessel surface (orthogonally) projected onto the image plane. This effect underestimates the actual area. The effect is stronger for endfeet close to the vessel “boundary” (vessel outline in the 2D image; in 3D there is no such boundary).
2. (Visibility error) The image only shows the part of the endfoot on the visible half of the vessel. An endfoot wrapping around the vessel will therefore once more be underestimated in area. Importantly, both discussed effects depend on the ratio 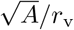, i.e. the endfoot size over the vessel radius. The effects are more pronounced for capillaries.

Consider the following two limit cases.

a. 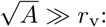 A small endfoot on a large vessel surface centered in the middle between the vessel boundaries is fully visible. There is only error (1). Since for an infinitely larger radius (limit case), this error tends to 0 for large vessels.
b. 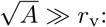 Imagine a rectangular endfoot (one side with arbitrary length) fully wrapped around the vessel (other side equal to 2*πr*_v_). Due to the projection error, the area is underestimated by a factor 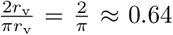. Additionally, only half of the endfoot is visible (factor 0.5). In combination, the area is underestimated by a factor 0.32, or the endfoot appears (on the image) approximately 3 times smaller than it actually is.

The presented theoretical model based on Voronoi tessellations of the vessel surface, allows us to investigate the combined error more systematically. To this end, we make one additional assumption. During the image analysis, seeing an image like Fig. 1 (bottom), the scientist is likely aware that counting small polygons close to the vessel boundary (corresponding to incompletely seen endfoot processes) decreases the accuracy of the results. Here, we cannot be sure how many such polygons have been counted. We therefore consider, in the theoretical analysis, several thresholds based on the centroid of the endfoot. Specifically, we assume that an endfoot is only counted if its centroid (w.r.t. to its visible projected portion) is in the middle *P* % between the vessel boundary. All polygons centered too close to the boundary on any side (somewhere in the (100 − *P*) % percent boundary region) will be omitted from the analysis. Since the value of *P* has a large influence on the results, we tested different numbers, (100 − *P*) ∈ {0, 10, 30, 50, 70}.

Next, we evaluated for different ratios 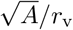 and different values of (100−*P*), the factor of endfoot area underestimation. To this end we generated patterns like in Fig. 1 (top), wrap them around the vessel, cf. Fig. 1 (bottom), and projected them onto the image plane. We did so from different angles and averages. The standard deviation of the computed factors was approximately 0.02 for all cases and is omitted in the following figure only showing the mean value. We averaged each parameter combination over 5 random realizations and report the mean value. The results are presented in Fig. 8. The results show that there is significant underestimation inherent to the projection, in particular for small vessels. For a capillary, with *r*_v_ = 3 µm and *A* = 50 µm, projection results in an area estimate between 15 µm and 35 µm. The upper value is obtained, if only polygons are counted that are centered in the middle 30 % of the seen vessel section. That means, even if the best is tried to exclude polygons that area clearly cut at the boundary, the error cannot be avoided.

**Figure 8.**
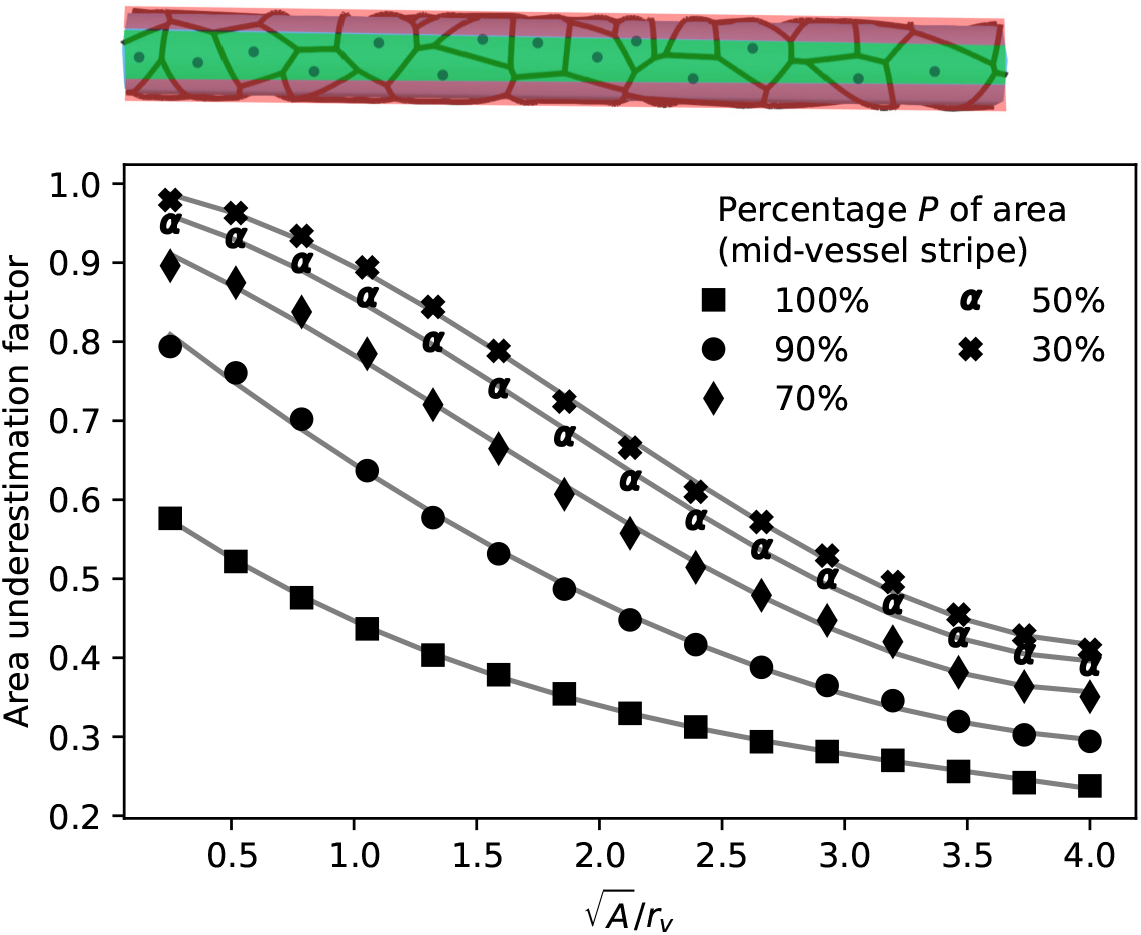
Evaluation of 2D image analysis error for endfoot area estimates. An example of the region of interest for one particular realization is shown in the top image. *P* = 100 means all polygons in the image have been counted. *P* = 30 means only the polygons centered in the stripe of thickness 0.3*d*_v_ in the middle of the vessel between the vessel outlines have been considered in the analysis. The top image shows *P* = 50 applied to the sample of Fig. 1 and the counted polygons are marked with a dot in their centroid. Counting all polygons leads to the most severe underestimation. The practice of discarding polygons close to the border reduces the error of the image analysis (underestimation fraction closer to 1). Solid lines are least squares curve fits with cubic polynomials.

Our analysis suggests a way to correct the error. To simplify the inverse problem of estimating the corrected area from the measured area, we choose *P* = 50 and observe that the least square fit with a cubic polynomial shown in Fig. 8 multiplied with the real area, 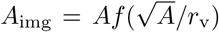, is monotone in the relevant parameter ranges (*r*_v_ ∈ [2.5, 40] µm, *A* ∈ [10, 500] µm^2^) and can therefore uniquely be inverted. Inverting the relationship *r*_v_ and *A*_img_ obtained by [6] at discrete sampling point with Brent’s root finding algorithm leads to the corrected diameter-area function shown in Fig. 3.

## B Pressure distribution in microvascular networks

The computed pressure distribution in microvascular networks used to classify vessels into arterial and venous vessels based on the network geometries extracted from measurement data [19] and boundary conditions estimated in [30, 28] are shown in Fig. 9.

**Figure 9.**
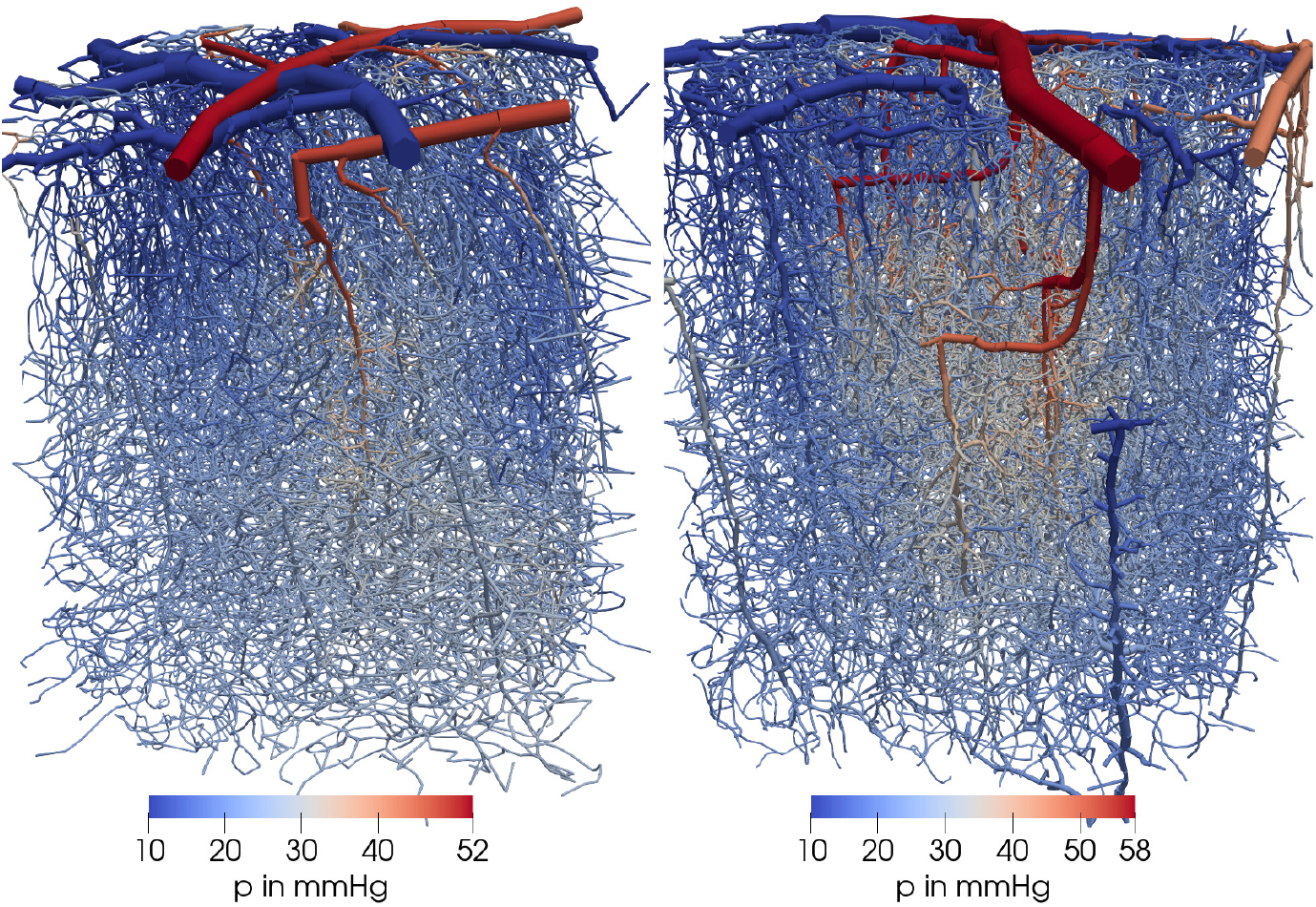
Two microvascular networks (MVN1/left, MVN2/right) of rodent cortex. Visualized in color is the computed blood pressure distribution. The vessels are shown as tubes scaled with the vessel lumen radius. Geometry based on data from [19].

## Ethics approval and consent to participate

Not applicable

## Consent for publication

Not applicable

## Availability of data and materials

All data generated or analyzed during this study are included in this published article. Any third-party data used is publicly available from the cited sources. The source code for the program used to generate Voronoi tessellations and the source code for the program used to compute pressure distributions in the microvascular networks in the current study are available from the corresponding author on request.

## Author’s contributions

TK conceived and designed the study, performed the simulations, analyzed the results, and wrote the article with input from the other authors; VV and KAM helped interpret the results and provide context. All authors read and approved the final manuscript.

## Acknowledgements

We would like to thank Franca Schmid for her support in interpreting the microvascular network data.

## List of abbreviations

CSF: cerebrospinal fluid;
ECS: extra-cellular space;
MVN: microvascular network;
MRI: magnetic resonance imaging;
MS: multiple sclerosis;
PVS: perivascular space.

## Footnotes

[1] This is to match the distribution of endfoot areas on single vessels measured by [6]. For example, a regular uniform cell center pattern would lead to uniform endfoot areas instead.

[2] using the open-source software CGAL [20]

[3] The endfoot processes forming the endfoot sheath overlap. The tessellation models the “visible” endfoot sheath surface configuration as seen from outside the vessel, cf. [6, Fig.2].

[4] *L* is chosen large (here *L* = 20*r*_o_) such that a possible bias on cell size due to boundary effects is minimized. The center points are duplicated along the sides so that the generated pattern is periodic and can be mapped onto (wrapped around) a cylinder surface (as shown in Fig. 1).

[5] Well-modeled based on a two-sample Kolmogorov-Smirnov test for goodness of fit. We remark that it has previously been observed that the cell area in Voronoi tessellations (with uniformly-distributed center point coordinates) may be approximated by a Gamma distribution, e.g. [36, 37, 38]; Weaire et al. [37] provide an intuitive explanation.

[6] computed as 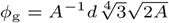 based on a single regular hexagon with area *A*

[7] For this estimate, the reported tissue permeabilities (3.5-14 × 10^−18^m^2^ [50], 0.4-110 × 10^−18^ m^2^ (mean: 16 × 10^−18^ ± 24 × 10^−18^ m^2^) [51], 10-20 × 10^−18^ m^2^ [52]) are divided by the slab thickness of 1 µm and fluid viscosity of 0.69 × 10^−3^ Pa s. We used a density of 1 × 10^3^ kg m^−3^ and gravitational acceleration of 9.81 m s^−1^ for conversion from hydraulic conductivity.

[8] In [53], the authors estimate tissue permeability based on data from (whole brain) convection-enhanced delivery studies, that are almost three orders of magnitude larger than what is reported from perfusion studies [50] and flow simulations [52]. However, the estimates might be altered in comparison with values for only ECS by effects of tissue deformation during injection [54] or by the inclusion or opening of highly permeable perivascular pathways.

[9] with Σ*SL*_*A*_ ≌ 1*/R*_IEG_ of [18] and *V* = 1 L.

[10] We mention that all three assumptions have been challenged and the current evidence does not suffice for a resolution of the debate, see e.g. [64] for a review. ^[11]^Large penetrating arterioles only account for about 1 % of the total microvascular surface area

## References

1. Wolff, J.: Beiträge zur Ultrastruktur der Kapillaren in der normalen Grosshirnrinde. Zeitschrift für Zellforschung und Mikroskopische Anatomie 60(3), 409–431 (1963). doi:10.1007/bf00336616

2. Helmchen, F., Kleinfeld, D.: In vivo measurements of blood flow and glial cell function with two-photon laser-scanning microscopy. In: Angiogenesis: In Vivo Systems, Part A. Methods in Enzymology, vol. 444, pp. 231–254. Academic Press, ??? (2008). Chap. 10. doi:10.1016/S0076-6879(08)02810-3

3. Mathiisen, T.M., Lehre, K.P., Danbolt, N.C., Ottersen, O.P.: The perivascular astroglial sheath provides a complete covering of the brain microvessels: An electron microscopic 3d reconstruction. Glia 58(9), 1094–1103 (2010). doi:10.1002/glia.20990

4. Watanabe, K., Takeishi, H., Hayakawa, T., Sasaki, H.: Three-dimensional organization of the perivascular glial limiting membrane and its relationship with the vasculature: A scanning electron microscope study. Okajimas Folia Anatomica Japonica 87(3), 109–121 (2010). doi:10.2535/ofaj.87.109

5. McCaslin, A.F.H., Chen, B.R., Radosevich, A.J., Cauli, B., Hillman, E.M.C.: In vivo 3d morphology of astrocyte—vasculature interactions in the somatosensory cortex: Implications for neurovascular coupling. Journal of Cerebral Blood Flow & Metabolism 31(3), 795–806 (2010). doi:10.1038/jcbfm.2010.204

6. Wang, M.X., Ray, L., Tanaka, K.F., Iliff, J.J., Heys, J.: Varying perivascular astroglial endfoot dimensions along the vascular tree maintain perivascular-interstitial flux through the cortical mantle. Glia 69(3), 715–728 (2020). doi:10.1002/glia.23923

7. Matsumae, M., Sato, O., Hirayama, A., Hayashi, N., Takizawa, K., Atsumi, H., Sorimachi, T.: Research into the physiology of cerebrospinal fluid reaches a new horizon: intimate exchange between cerebrospinal fluid and interstitial fluid may contribute to maintenance of homeostasis in the central nervous system. Neurologia medico-chirurgica 56(7), 416–441 (2016)

8. Iliff, J.J., Wang, M., Liao, Y., Plogg, B.A., Peng, W., Gundersen, G.A., Benveniste, H., Vates, G.E., Deane, R., Goldman, S.A., Nagelhus, E.A., Nedergaard, M.: A paravascular pathway facilitates CSF flow through the brain parenchyma and the clearance of interstitial solutes, including amyloid β. Science Translational Medicine 4(147) (2012). doi:10.1126/scitranslmed.3003748

9. Asgari, M., de Zélicourt, D., Kurtcuoglu, V.: How astrocyte networks may contribute to cerebral metabolite clearance. Scientific Reports 5(1) (2015). doi:10.1038/srep15024

10. MacAulay, N.: Molecular mechanisms of brain water transport. Nature Reviews Neuroscience 22(6), 326–344 (2021). doi:10.1038/s41583-021-00454-8

11. MacAulay, N.: Reply to ‘aquaporin 4 and glymphatic flow have central roles in brain fluid homeostasis’. Nature Reviews Neuroscience 22(10), 651–652 (2021). doi:10.1038/s41583-021-00515-y

12. Honda, H.: Description of cellular patterns by dirichlet domains: The two-dimensional case. Journal of Theoretical Biology 72(3), 523–543 (1978). doi:10.1016/0022-5193(78)90315-6

13. Zisis, E., Keller, D., Kanari, L., Arnaudon, A., Gevaert, M., Delemontex, T., Coste, B., Foni, A., Abdellah, M., Calì, C., Hess, K., Magistretti, P.J., Schürmann, F., Markram, H.: Digital reconstruction of the neuro-glia-vascular architecture. Cerebral Cortex 31(12), 5686–5703 (2021). doi:10.1093/cercor/bhab254

14. Brightman, M.W., Reese, T.S.: Junctions between intimately apposed cell membranes in the vertebrate brain. Journal of Cell Biology 40(3), 648–677 (1969). doi:10.1083/jcb.40.3.648

15. Koch, T., Flemisch, B., Helmig, R., Wiest, R., Obrist, D.: A multiscale subvoxel perfusion model to estimate diffusive capillary wall conductivity in multiple sclerosis lesions from perfusion MRI data. International Journal for Numerical Methods in Biomedical Engineering 36(2) (2020). doi:10.1002/cnm.3298

16. Tithof, J., Boster, K.A., Bork, P.A., Nedergaard, M., Thomas, J.H., Kelley, D.H.: A network model of glymphatic flow under different experimentally-motivated parametric scenarios. iScience 25(5), 104258 (2022). doi:10.1016/j.isci.2022.104258

17. Faghih, M.M., Sharp, M.K.: Is bulk flow plausible in perivascular, paravascular and paravenous channels? Fluids and Barriers of the CNS 15(1), 1–10 (2018). doi:10.1186/s12987-018-0103-8

18. Vinje, V., Eklund, A., Mardal, K.-A., Rognes, M.E., Støverud, K.-H.: Intracranial pressure elevation alters CSF clearance pathways. Fluids and Barriers of the CNS 17(1) (2020). doi:10.1186/s12987-020-00189-1

19. Blinder, P., Tsai, P.S., Kaufhold, J.P., Knutsen, P.M., Suhl, H., Kleinfeld, D.: The cortical angiome: an interconnected vascular network with noncolumnar patterns of blood flow. Nature Neuroscience 16(7), 889–897 (2013). doi:10.1038/nn.3426

20. Yvinec, M.: 2D triangulations. In: CGAL User and Reference Manual, 5.5 edn. CGAL Editorial Board, ??? (2022). https://doc.cgal.org/5.5/Manual/packages.html#PkgTriangulation2

21. Rohatgi, A.: Webplotdigitizer: Version 4.5 (2021). https://automeris.io/WebPlotDigitizer

22. Nicholson, C., Hrabětová, S.: Brain extracellular space: The final frontier of neuroscience. Biophysical Journal 113(10), 2133–2142 (2017). doi:10.1016/j.bpj.2017.06.052

23. Michel, C.C., Curry, F.E.: Microvascular permeability. Physiological Reviews 79(3), 703–761 (1999). doi:10.1152/physrev.1999.79.3.703

24. Renkin, E.M.: Filtration, diffusion, and molecular sieving through porous cellulose membranes. The Journal of general physiology 38(2), 225 (1954)

25. Beck, R.E., Schultz, J.S.: Hindrance of solute diffusion within membranes as measured with microporous membranes of known pore geometry. Biochimica et Biophysica Acta (BBA) - Biomembranes 255(1), 273–303 (1972). doi:10.1016/0005-2736(72)90028-4

26. Herring, N., Paterson, D.J.: Levick’s Introduction to Cardiovascular Physiology, Sixth Edition. CRC Press, ??? (2018). doi:10.1201/9781351107754

27. Thomas, J.A., McGaughey, A.J.H., Kuter-Arnebeck, O.: Pressure-driven water flow through carbon nanotubes: Insights from molecular dynamics simulation. International Journal of Thermal Sciences 49(2), 281–289 (2010). doi:10.1016/j.ijthermalsci.2009.07.008

28. Schmid, F.: Averaged results of blood flow simulations with discrete RBC tracking for microvascular networks. Zenodo (2017). doi:10.5281/zenodo.269650

29. Douglas, D.H., Peucker, T.K.: Algorithms for the reduction of the number of points required to represent a digitized line or its caricature. Cartographica: The International Journal for Geographic Information and Geovisualization 10(2), 112–122 (1973). doi:10.3138/fm57-6770-u75u-7727

30. Schmid, F., Tsai, P.S., Kleinfeld, D., Jenny, P., Weber, B.: Depth-dependent flow and pressure characteristics in cortical microvascular networks. PLOS Computational Biology 13(2), 1005392 (2017). doi:10.1371/journal.pcbi.1005392

31. Pries, A.R., Secomb, T.W., Gessner, T., Sperandio, M.B., Gross, J.F., Gaehtgens, P.: Resistance to blood flow in microvessels in vivo. Circulation Research 75(5), 904–915 (1994). doi:10.1161/01.res.75.5.904

32. Koch, T., Gläser, D., Weishaupt, K., et al.: DuMux 3 – an open-source simulator for solving flow and transport problems in porous media with a focus on model coupling. Computers & Mathematics with Applications 81, 423–443 (2021). doi:10.1016/j.camwa.2020.02.012

33. Sander, O., Koch, T., Schröder, N., Flemisch, B.: The dune foamgrid implementation for surface and network grids. Archive of Numerical Software Vol 5, 1–2017 (2017). doi:10.11588/ANS.2017.1.28490

34. Hill, R.A., Tong, L., Yuan, P., Murikinati, S., Gupta, S., Grutzendler, J.: Regional blood flow in the normal and ischemic brain is controlled by arteriolar smooth muscle cell contractility and not by capillary pericytes. Neuron 87(1), 95–110 (2015). doi:10.1016/j.neuron.2015.06.001

35. Bonney, S.K., Coelho-Santos, V., Huang, S.-F., Takeno, M., Kornfeld, J., Keller, A., Shih, A.Y.: Public volume electron microscopy data: An essential resource to study the brain microvasculature. Frontiers in Cell and Developmental Biology 10 (2022). doi:10.3389/fcell.2022.849469

36. Kiang, T.: Random Fragmentation in Two and Three Dimensions. Zeitschrift für Astrophysik 64, 433 (1966)

37. Weaire, D., Kermode, J.P., Wejchert, J.: On the distribution of cell areas in a voronoi network. Philosophical Magazine B 53(5), 101–105 (1986). doi:10.1080/13642818608240647

38. Koufos, K., Dettmann, C.P.: Distribution of cell area in bounded poisson voronoi tessellations with application to secure local connectivity. Journal of Statistical Physics 176(5), 1296–1315 (2019). doi:10.1007/s10955-019-02343-y

39. Korogod, N., Petersen, C.C., Knott, G.W.: Ultrastructural analysis of adult mouse neocortex comparing aldehyde perfusion with cryo fixation. eLife 4 (2015). doi:10.7554/elife.05793

40. Haj-Yasein, N.N., Vindedal, G.F., Eilert-Olsen, M., Gundersen, G.A., Øivind Skare, Laake, P., Klungland, A., Thorén, A.E., Burkhardt, J.M., Ottersen, O.P., Nagelhus, E.A.: Glial-conditional deletion of aquaporin-4 (Aqp4) reduces blood-brain water uptake and confers barrier function on perivascular astrocyte endfeet. Proceedings of the National Academy of Sciences 108(43), 17815–17820 (2011). doi:10.1073/pnas.1110655108

41. Kubotera, H., Ikeshima-Kataoka, H., Hatashita, Y., Allegra Mascaro, A.L., Pavone, F.S., Inoue, T.: Astrocytic endfeet re-cover blood vessels after removal by laser ablation. Scientific Reports 9(1), 1263 (2019). doi:10.1038/s41598-018-37419-4

42. Mills, W.A., Woo, A.M., Jiang, S., Martin, J., Surendran, D., Bergstresser, M., Kimbrough, I.F., Eyo, U.B., Sofroniew, M.V., Sontheimer, H.: Astrocyte plasticity in mice ensures continued endfoot coverage of cerebral blood vessels following injury and declines with age. Nature Communications 13(1) (2022). doi:10.1038/s41467-022-29475-2

43. Florence, C.M., Baillie, L.D., Mulligan, S.J.: Dynamic volume changes in astrocytes are an intrinsic phenomenon mediated by bicarbonate ion flux. PLoS ONE 7(11), 51124 (2012). doi:10.1371/journal.pone.0051124

44. Rosic, A.B., Dukefoss, D.B., Åbjørsbråten, K.S., Tang, W., Jensen, V., Ottersen, O.P., Enger, R., Nagelhus, E.A.: Aquaporin-4-independent volume dynamics of astroglial endfeet during cortical spreading depression. Glia 67(6), 1113–1121 (2019). doi:10.1002/glia.23604

45. Schmid, F., Barrett, M.J.P., Jenny, P., Weber, B.: Vascular density and distribution in neocortex. NeuroImage 197, 792–805 (2019). doi:10.1016/j.neuroimage.2017.06.046

46. Bojarskaite, L., Bjørnstad, D.M., Vallet, A., Binder, K.M.G., Cunen, C., Heuser, K., Kuchta, M., Mardal, K.-A., Enger, R.: Sleep cycle-dependent vascular dynamics enhance perivascular cerebrospinal fluid flow and solute transport. BioRxiv preprint (2022). doi:10.1101/2022.07.14.500017

47. Syková, E., Nicholson, C.: Diffusion in brain extracellular space. Physiological Reviews 88(4), 1277–1340 (2008). doi:10.1152/physrev.00027.2007

48. Kimura, M., Dietrich, H.H., Huxley, V.H., Reichner, D.R., Dacey, R.G.: Measurement of hydraulic conductivity in isolated arterioles of rat brain cortex. American Journal of Physiology-Heart and Circulatory Physiology 264(6), 1788–1797 (1993). doi:10.1152/ajpheart.1993.264.6.h1788

49. Fraser, P.A., Dallas, A.D., Davies, S.: Measurement of filtration coefficient in single cerebral microvessels of the frog. The Journal of Physiology 423(1), 343–361 (1990). doi:10.1113/jphysiol.1990.sp018026

50. Swabb, E.A., Wei, J., Gullino, P.M.: Diffusion and convection in normal and neoplastic tissues. Cancer Research 34(10), 2814–2822 (1974)

51. Franceschini, G., Bigoni, D., Regitnig, P., Holzapfel, G.A.: Brain tissue deforms similarly to filled elastomers and follows consolidation theory. Journal of the Mechanics and Physics of Solids 54(12), 2592–2620 (2006). doi:10.1016/j.jmps.2006.05.004

52. Holter, K.E., Kehlet, B., Devor, A., Sejnowski, T.J., Dale, A.M., Omholt, S.W., Ottersen, O.P., Nagelhus, E.A., Mardal, K.-A., Pettersen, K.H.: Interstitial solute transport in 3d reconstructed neuropil occurs by diffusion rather than bulk flow. Proceedings of the National Academy of Sciences 114(37), 9894–9899 (2017). doi:10.1073/pnas.1706942114

53. Smith, J.H., Humphrey, J.A.C.: Interstitial transport and transvascular fluid exchange during infusion into brain and tumor tissue. Microvascular Research 73(1), 58–73 (2007). doi:10.1016/j.mvr.2006.07.001

54. Støverud, K.H., Darcis, M., Helmig, R., Hassanizadeh, S.M.: Modeling concentration distribution and deformation during convection-enhanced drug delivery into brain tissue. Transport in porous media 92(1), 119–143 (2012)

55. Vidotto, E., Koch, T., Köppl, T., Helmig, R., Wohlmuth, B.: Hybrid models for simulating blood flow in microvascular networks. Multiscale Modeling & Simulation 17(3), 1076–1102 (2019). doi:10.1137/18m1228712

56. Keller, D., Erö, C., Markram, H.: Cell densities in the mouse brain: A systematic review. Frontiers in Neuroanatomy 12 (2018). doi:10.3389/fnana.2018.00083

57. Ji, X., Ferreira, T., Friedman, B., Liu, R., Liechty, H., Bas, E., Chandrashekar, J., Kleinfeld, D.: Brain microvasculature has a common topology with local differences in geometry that match metabolic load. Neuron 109(7), 1168–118713 (2021). doi:10.1016/j.neuron.2021.02.006

58. Adams, D.L., Piserchia, V., Economides, J.R., Horton, J.C.: Vascular supply of the cerebral cortex is specialized for cell layers but not columns. Cerebral Cortex 25(10), 3673–3681 (2014). doi:10.1093/cercor/bhu221

59. Iliff, J.J., Wang, M., Liao, Y., Plogg, B.A., Peng, W., Gundersen, G.A., Benveniste, H., Vates, G.E., Deane, R., Goldman, S.A., Nagelhus, E.A., Nedergaard, M.: A paravascular pathway facilitates CSF flow through the brain parenchyma and the clearance of interstitial solutes, including amyloid β. Science Translational Medicine 4(147) (2012). doi:10.1126/scitranslmed.3003748

60. Mestre, H., Tithof, J., Du, T., Song, W., Peng, W., Sweeney, A.M., Olveda, G., Thomas, J.H., Nedergaard, M., Kelley, D.H.: Flow of cerebrospinal fluid is driven by arterial pulsations and is reduced in hypertension. Nature communications 9(1), 1–9 (2018). doi:10.1038/s41467-018-07318-3

61. Eide, P.K., Valnes, L.M., Lindstrøm, E.K., Mardal, K.-A., Ringstad, G.: Direction and magnitude of cerebrospinal fluid flow vary substantially across central nervous system diseases. Fluids and Barriers of the CNS 18(1), 1–18 (2021). doi:10.1186/s12987-021-00251-6

62. Eide, P.K., Vinje, V., Pripp, A.H., Mardal, K.-A., Ringstad, G.: Sleep deprivation impairs molecular clearance from the human brain. Brain 144(3), 863–874 (2021). doi:10.1093/brain/awaa443

63. Xie, L., Kang, H., Xu, Q., Chen, M.J., Liao, Y., Thiyagarajan, M., O’Donnell, J., Christensen, D.J., Nicholson, C., Iliff, J.J., et al.: Sleep drives metabolite clearance from the adult brain. science 342(6156), 373–377 (2013). doi:10.1126/science.1241224

64. Hladky, S.B., Barrand, M.A.: The glymphatic hypothesis: the theory and the evidence. Fluids and Barriers of the CNS 19(1) (2022). doi:10.1186/s12987-021-00282-z

